# Multivariate mining of an alpaca immune repertoire identifies potent cross-neutralising SARS-CoV-2 nanobodies

**DOI:** 10.1101/2021.07.25.453673

**Authors:** Leo Hanke, Daniel J. Sheward, Alec Pankow, Laura Perez Vidakovics, Vivien Karl, Changil Kim, Egon Urgard, Natalie L. Smith, Juan Astorga-Wells, Simon Ekström, Jonathan M. Coquet, Gerald M. McInerney, Ben Murrell

## Abstract

Conventional approaches to isolate and characterize nanobodies are laborious and cumbersome. Here we combine phage display, multivariate enrichment, and novel sequence analysis techniques to annotate an entire nanobody repertoire from an immunized alpaca. We combine this approach with a streamlined screening strategy to identify numerous anti-SARS-CoV-2 nanobodies, and use neutralization assays and Hydrogen/Deuterium exchange coupled to mass spectrometry (HDX-MS) epitope mapping to characterize their potency and specificity. Epitope mapping revealed that the binding site is a key determinant of neutralization potency, rather than affinity alone. The most potent nanobodies bind to the receptor binding motif of the RBD, directly preventing interaction with the host cell receptor ACE2, and we identify two exceptionally potent members of this category (with monomeric IC50s around 13 and 16 ng/ml). Other nanobodies bind to a more conserved epitope on the side of the RBD, and are able to potently neutralize the SARS-CoV-2 founder virus (42 ng/ml), the beta variant (B.1.351/501Y.V2) (35 ng/ml), and also cross-neutralize the more distantly related SARS-CoV-1 (0.46 μg/ml). The approach presented here is well suited for the screening of phage libraries to identify functional nanobodies for various biomedical and biochemical applications.

## Introduction

Camelids, including llamas and alpacas, express unique immunoglobulins composed of just heavy chains ^1^. The antigen-binding variable fragment is a single-domain that can be expressed recombinantly as a 15 kDa antibody fragment called VHH or nanobody. For applications where the functions of an Fc-domain are not required, nanobodies have many advantages over their full-size antibody counterparts. Nanobodies can be produced at high quantities much more cost-effectively than monoclonal antibodies. Their small size and single-gene nature allows for easy cloning, modification, and functionalization and also permits better tissue penetration and, for imaging purposes, closer proximity of fluorophores or radioisotopes to the antigen. As a result, nanobodies have applications in cell biology ^2^, structural biology ^3^, cancer research ^4^, and immunology ^5^. In addition, nanobodies are ideal neutralizing molecules or perturbants of viruses. Potent SARS-CoV-2 neutralizing nanobodies target the receptor binding domain (RBD) of the spike protein. They neutralize the virus either by blocking ACE2 receptor interactions ^6,7^, or other mechanisms such as the triggering of conformational changes ^8^. One affinity matured nanobody fused to a human IgG is currently in clinical development for treatment of SARS-CoV-2 infections ^9^.

Antigen-specific nanobodies are typically isolated from large immune libraries ^10,11^, or more recently, also from synthetic libraries ^12,13^. Typically, such libraries are screened using robust phage display or yeast display techniques. To encompass the larger diversity of non-immune and synthetic libraries, screening typically starts with ribosome display. Often these screens only yield a handful of useful binders. Other screens using lentiviral nanobody libraries allow direct phenotypic readouts that are more productive and limit the time-consuming functional testing of individual identified binders ^14^. However, phenotypic readouts cannot be implemented in all cases. To address this gap, we combined the robust and versatile phage-display with rapid high-throughput functional testing, both bridged by next-generation sequencing (NGS) and enrichment analysis. Applying this approach, we identify a panel of potent SARS-CoV-2 neutralizing nanobodies, and provide detailed methodological descriptions enabling easy implementation in other nanobody discovery workflows.

## Results

To generate a library of nanobodies against SARS-CoV-2, we immunized one alpaca four times with prefusion-stabilized SARS-CoV-2 spike protein and the RBD (Fig 1). For each immunization, both proteins were injected separately into different flanks of the animal. Four days after the last immunization, we isolated peripheral blood mononuclear cells (PBMCs), amplified nanobody specific regions, and constructed a phagemid library. We performed three parallel phage selections on proteins C-terminally immobilized on magnetic beads. The first screen was performed with recombinant spike protein (“S”), the second screen with RBD (“RBD”), and the third screen with immobilized spike in the presence of non-immobilized RBD to deplete RBD-specific nanobodies from the phage pool (“ScRBD”). Thus, the RBD and ScRBD pannings should enrich for non-overlapping sets of nanobodies, and the S panning should enrich for the union of these.

**Fig. 1.**
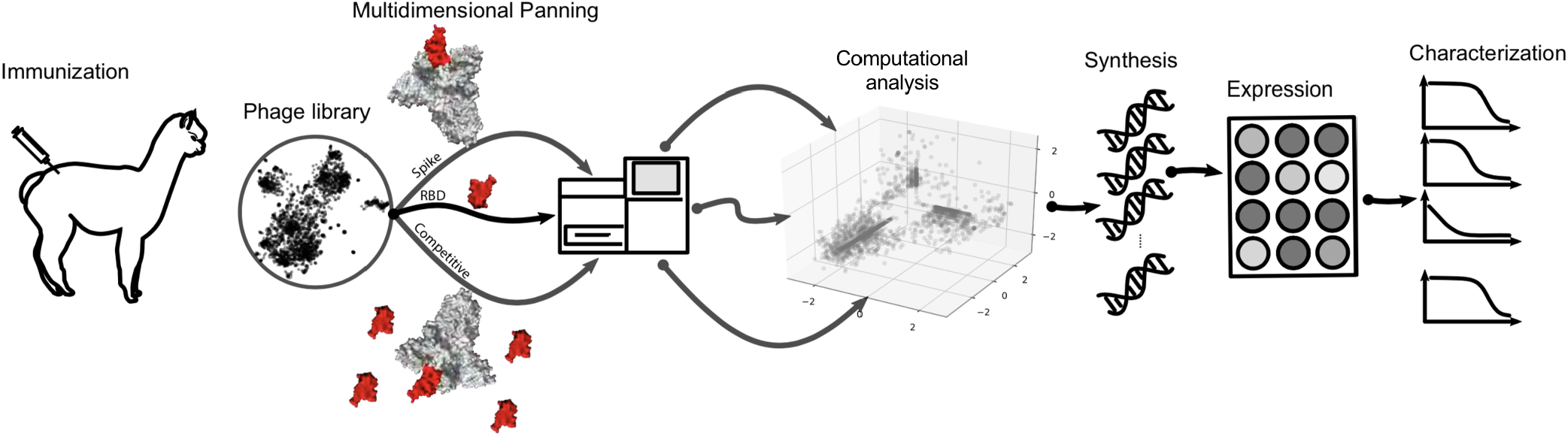
Strategy overview. A nanobody phage library is constructed from the immune repertoire of an immunized alpaca. This undergoes multiple independent panning steps, each enriching for distinct epitope targets. The original library and the enriched population from each panning are deeply sequenced. Computational enrichment analysis characterizes the repertoire, and aids in the selection of nanobody variants for synthesis, expression, and downstream characterization.

### Enrichment analysis by NGS

Phage pools were sequenced by Illumina, both prior to enrichment, as well as after each panning step. The premise of our approach was that when the starting library contains a large number of distinct variants with highly variable frequencies (as is expected of immune repertoires ^15^), the final frequency post-panning is dominated by the starting frequency of a variant. The enrichment - the increase in frequency due to panning - which should be a better proxy for binding affinity, can be overwhelmed by the starting frequency. This suggests that the traditional approach of picking colonies post-panning ^11^ may miss potent nanobodies with low starting frequencies. We calculate the enrichment for each variant as the log ratio of the frequency post-panning over pre-panning, regularized with a pseudocount ^16^ (to accommodate variants that are only observed after panning). To base our choices on enrichment, we need to know how reliable are the estimates of enrichment. Here we exploit the fact that the library construction step includes a primer with degenerate bases, allowing us to consider independent versions of each variant that differ at a synonymous position. A correlation plot of the enrichment for two versions of each variant shows excellent agreement for RBD and ScRBD, but weak agreement for S (See SI fig 1). Possible explanations for this reduced correlation could be due to S offering more targets to the nanobody repertoire, increasing the competition and perhaps the stochasticity, or the introduction of a bottleneck at the start of the panning step, causing some variants to drop out. Regardless of the explanation, this indicates that enrichment calculated from the S panning may provide a less reliable signal of enrichment than RBD and ScRBD pannings. For further analysis, the cloning primer regions were ignored to provide total counts for all versions of each variant, from which the final enrichment metrics were calculated. Figure 2A shows the RBD against the ScRBD enrichment. As intended by the panning design, there were very few sequence variants showing enrichment in both of these panning steps. Further, we color by S enrichment, which tends to be higher when either RBD or ScRBD enrichment is high, but the unreliability of the S enrichment identified by the barcode analysis is also visible at this level, especially for variants that were smaller in the baseline library.

**Fig. 2.**
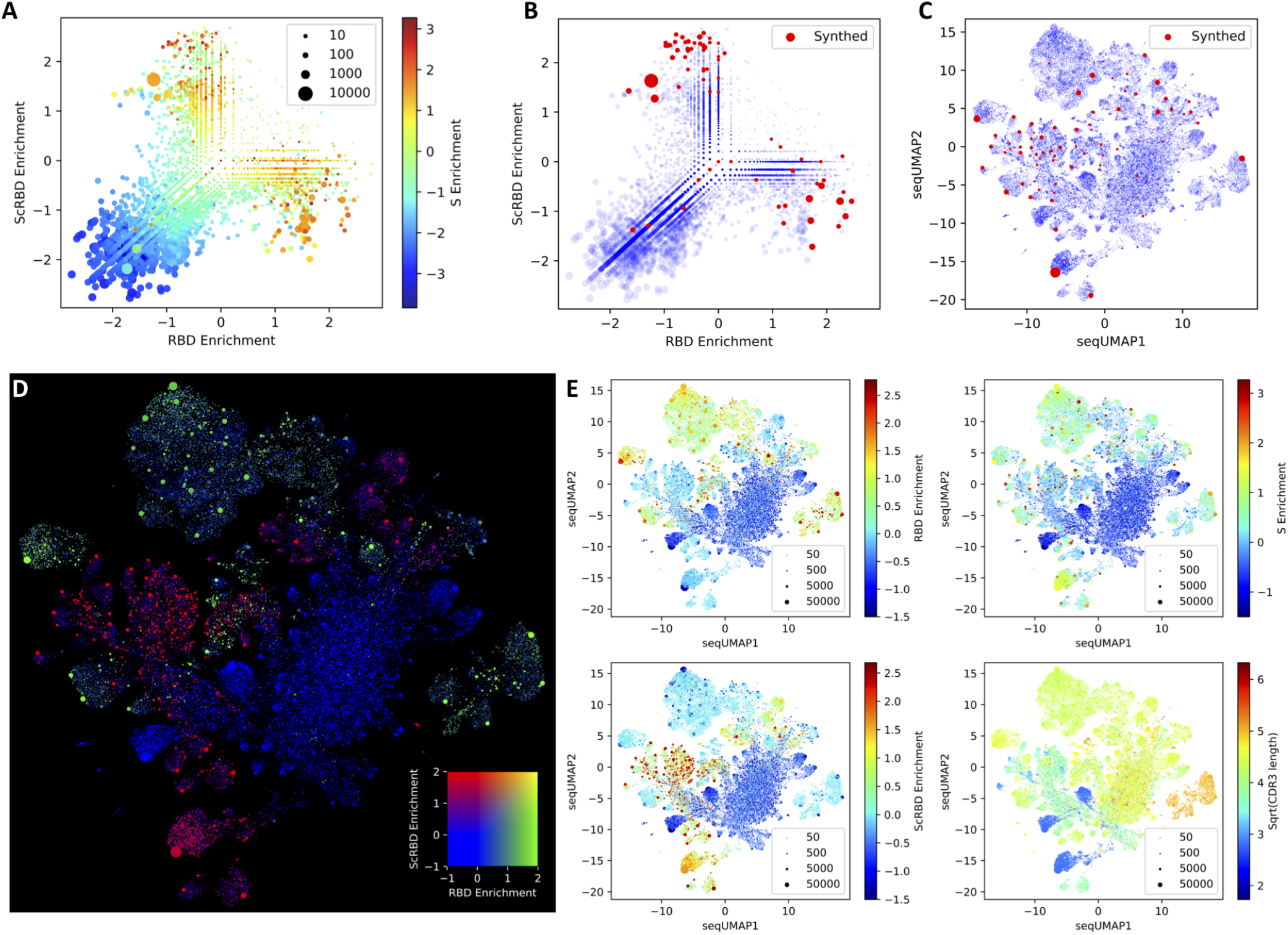
Multivariate repertoire analysis. A shows the enrichment across three parallel panning runs, with RBD panning on the x axis, spike with competing soluble RBD (“ScRBD”) on the y axis, and colored by spike (“S”). Variants selected for further screening are shown either upon the enrichment plot (B), or upon a seqUMAP embedding of the nanobody sequences (C), which embeds nanobody sequences into two dimensions such that closely related variants are neighbours. D shows which regions of seqUMAP space are targeting RBD (green), which are targeting the rest of spike (red), and which are not SARS-CoV-2 specific (blue). The lack of double-enrichment (yellow) in RBD and ScRBD shows that these two panning runs enriched for mutually exclusive variants, which is corroborated by the lack of points in the top right quadrant of panel A. E shows RBD, S, and ScRBD enrichment separately, as well as the CDR3 lengths (number of amino acids, square root transformed) for all nanobody variants overlaid on the seqUMAP plot.

### Visualizing nanobody repertoire VDJ space

When selecting nanobody variants, a key dimension is their relatedness. For screening purposes, nanobodies with similar VDJ sequences should be avoided, but later it might be useful to screen further candidates related to any promising hits. This would be aided by a way of visualizing sequence relatedness. One standard approach for visualizing a set of sequences would be a phylogeny, or clustering dendrogram, but these are unwieldy for such large sequence datasets, often requiring multiple sequence alignments, and behaving poorly for regions of problematic homology, which are especially common in nanobody CDR3s. Here we adapt the Uniform Manifold Approximation and Projection (UMAP ^17^) - an approach that is now standard in the single-cell RNAseq literature and popular in many other domains), but is not commonly used to visualize sequence data, possibly due to technical challenges. We circumvent these issues using kmer sequence embeddings ^18^, and call the resulting approach “seqUMAP”, which embeds sequences into two-dimensional space, such that closely related sequences are neighbours.

Figure 2C-E shows seqUMAP embeddings of the entire nanobody repertoire, overlaid with different data for each variant. Where enrichment is plotted (especially for RBD and ScRBD, but less so for S), there is a striking spatial association with enrichment, showing that genetic relatedness strongly predicts whether a variant is enriched. The mutual exclusivity of RBD and ScRBD panning is recapitulated in Figure 2D, where entire regions of VDJ space are enriched exclusively in one or the other, but not both. We selected 72 nanobodies from across enrichment and VDJ space, shown in Figure 2B and C. Since less is known about anti-spike nanobodies that do not target the RBD, we biased our selection to include approximately twice as many ScRBD enriched candidates as RBD enriched candidates.

### Enrichment Predicts Binding

The 72 selected nanobodies were synthesized and cloned into a nanobody expression vector. Nanobodies were expressed in a 96-deep-well plate with a culture volume of 1 ml. Expressed nanobodies were retrieved from the periplasm by osmotic shock, and the periplasmic extract was analyzed by SDS-PAGE and Coomassie staining. A band corresponding to the nanobody was visible for most periplasmic extracts at 15 kDa, alongside other bands typically of higher molecular weight (Fig. S3). To get an estimate of the expression efficiency of the different nanobodies, we quantified the band intensity. For most nanobodies, expression efficiency and purity were sufficient to analyze binding specificity by ELISA (spike or RBD coated) and antiviral activity by pseudotyped virus neutralization assays. Figure 3 shows the enrichment metrics from the NGS data as well as the ELISA values for all selected nanobodies. We pair the ScRBD enrichment with the difference between the Spike and RBD ELISA values. With only a few exceptions, nanobodies that were enriched for a particular target show ELISA signal for that target, with correlation coefficients of r=0.72 for RBD (p < 10^−5^) and r=0.66 for ScRBD (p < 10^−5^). The correlation for S was not significant, which is mostly because both spike and RBD targets exhibit S ELISA signal, reducing the variance, but may be due, in part, to the less reliable enrichment estimates for S than for RBD or ScRBD.

**Fig. 3.**
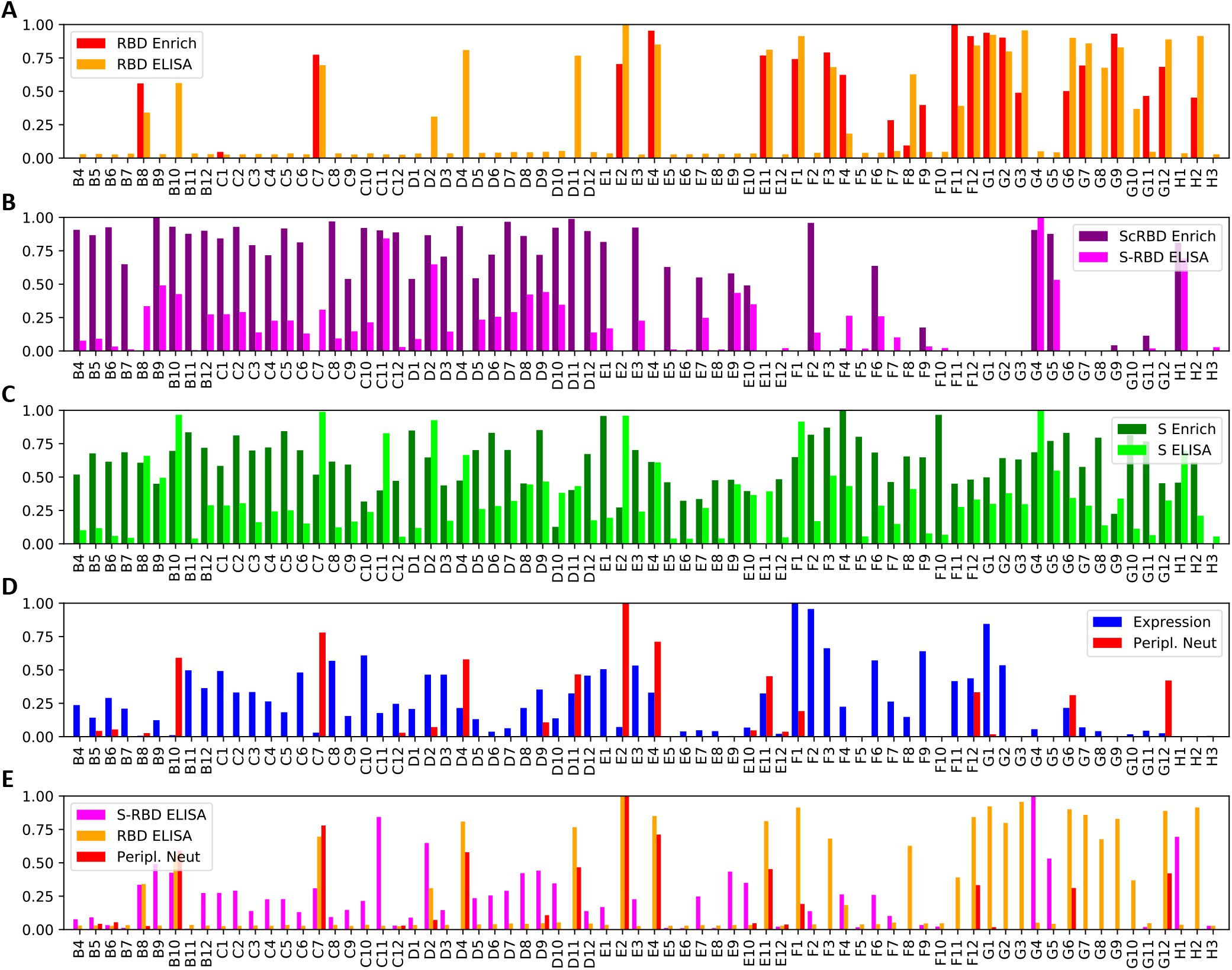
Rapid nanobody screening. 72 nanobodies, selected from the multivariate analysis, were synthesized and expressed, and the crude periplasmic extract screened for expression, binding, and neutralization. All values are normalized to the maximum value across nanobodies. The top three plots depict, for each nanobody, the enrichment calculated from the NGS data and the corresponding periplasmic extract ELISA. The ELISA signal in the second plot is the RBD OD450 subtracted from the S OD450, and should only be strongly positive when a nanobody binds spike outside of the RBD. The top two panels show that, for the vast majority of nanobody variants, the enrichment analysis is strongly predictive of whether the nanobody targets RBD or not. The bottom panel shows expression and (log-domain) pseudotyped virus neutralization IC50s, and nanobody expression calculated from the periplasmic extract.

### Neutralization

To identify nanobodies capable of neutralizing SARS-CoV-2 we employed a high-throughput pseudotyped-virus neutralization assay, directly assessing neutralization by periplasmic extracts over 4 serial 3-fold dilutions. We observed a common baseline signal for inhibition at low dilutions relative to wells without periplasmic extract, which we therefore subtracted from all measurements. This screen identified a number of nanobodies displaying potent neutralizing capacity. Normalized log pseudotyped virus neutralization titers are shown in figure 3DE, alongside normalized expression results and ELISA values. We selected 13 candidates for downstream analysis, including the most potently neutralizing RBD-specific nanobodies as well as several non-RBD binders, that were then expressed and purified. Neutralizing antibody titers of purified nanobodies were highly correlated with the preliminary periplasm screens, with nanobodies C7 and E2 displaying exceptional potency with IC50s in the range of 0.01 μg/ml (Fig 4A). Note that E4 is identical to the “Fu2” nanobody that was isolated via more traditional colony picking from the same immunized animal, and is extensively described elsewhere ^19^. Two of the selected non-RBD-specific nanobodies (C11, D9) were also capable of neutralization, albeit weakly and for D9 plateauing at approximately 50

**Fig. 4.**
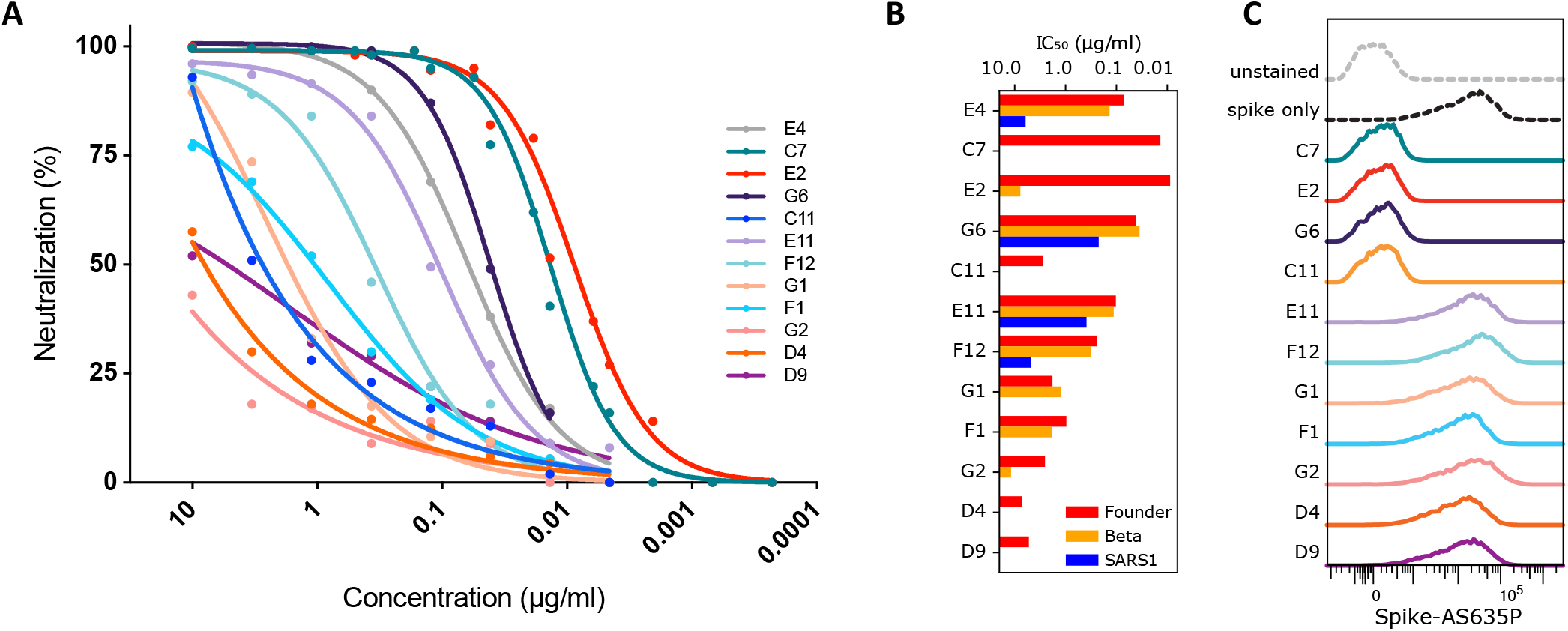
Neutralization by nanobody monomers. 13 candidate nanobodies, selected from the rapid screen, were profiled by pseudotyped lentivirus assay for neutralization activity. A depicts neutralization curves. B shows neutralization IC50s against the SARS-CoV-2 “Founder” variant (Wu-Hu-1), the Beta variant of concern (first described in South Africa - 501Y.V2/B.1.351), and the more distantly-related SARS-CoV-1 (SARS1). C depicts the intensity of labelled spike in a flow cytometry assay, showing whether a nanobody can prevent fluorescent spike protein from binding to HEK293T-hACE2 target cells, clarifying the mechanism of neutralization.

### A subset of nanobodies are broadly neutralizing

Variants of Concern ^20–22^ are rapidly rising in frequency. Some of these exhibit mutations that confer escape from prior immunity and from many existing monoclonal antibody therapy candidates ^23^. Given this context, one approach to addressing this problem is to attempt to discover broadly neutralizing biologics. Fig. 4B shows that many of the identified nanobodies are sufficiently broad to neutralize both the SARS-CoV-2 “founder” variant, and the beta Variant of Concern (B.1.351/501Y.V2), sometimes without any reduction in potency. The most potently neutralizing nanobodies, C7 and E2, lack any meaningful cross-neutralization, but G6 is exceptionally potent against beta (IC50 = 35 ng/ml). Furthermore, two nanobodies (E11 and G6) show substantial cross-neutralization of SARS-CoV-1, which is a far more distantly related member of the betacoronavirus genus, suggesting the targeting of a more conserved epitope.

### Nanobodies do not need to bind the RBD or block ACE2 receptor interaction to neutralize SARS-CoV-2

Nanobodies have been shown to neutralize SARS-CoV-2 by various mechanisms, including direct competition with ACE2^6,7^, locking of RBDs in an ACE2-inaccessible conformation ^24^, dimerizing spikes and agglutinating virions ^19^, or triggering the post-fusion conformation and shedding of S1^8^. To test if identified nanobodies interfere with the binding of the spike protein to the hACE2 receptor we performed a flow cytometry-based competitive binding assay. Human ACE2 (hACE2) expressing HEK293T cells were stained with fluorescently labeled prefusion-stabilized spike trimers, either alone, or preincubated with saturating amounts of the different nanobodies (Fig. 4C). Preincubation of fluorescently labeled spike protein with the most potent nanobodies, C7 and E2, completely abolished staining of the hACE2 positive cells. Similarly, G6 effectively prevented spike binding to ACE2. Surprisingly, one nanobody (C11) that binds to an non-RBD-epitope was still capable of preventing spike binding to ACE2 expressing cells. In contrast, other tested nanobodies did not reduce staining suggesting that they neutralize SARS-CoV-2 by mechanisms other than blocking ACE2 interaction.

### RBD-specific nanobodies bind with subnanomolar affinity

Binding affinities can impact neutralization potential. We determined the binding kinetics of five RBD-specific nanobodies by surface plasmon resonance (SPR). We observed high affinities in the picomolar range for all tested nanobodies (C7, E2, E11, F1, G6) (Fig. 5A). F1, despite only moderate neutralization capability, showed the highest affinity, with a barely detectable dissociation rate in our assay. We conclude that the kD of F1 to the RBD is < 10^−11^. For the nanobody C7, we noticed a poor fit of the 1:1 Langmuir model, and the elution profile during size-exclusion purification indicated the tendency for natural dimer- and multimerization. Indeed, the heterogeneous binding model fit the C7 SPR sensorgrams well, confirming that two binding events are observed simultaneously: binding of C7 to the RBD and binding of C7 to C7. Overall, for these five nanobodies, binding affinities do not straightforwardly correlate with neutralization, suggesting that other nanobody characteristics determine potency.

**Fig. 5.**
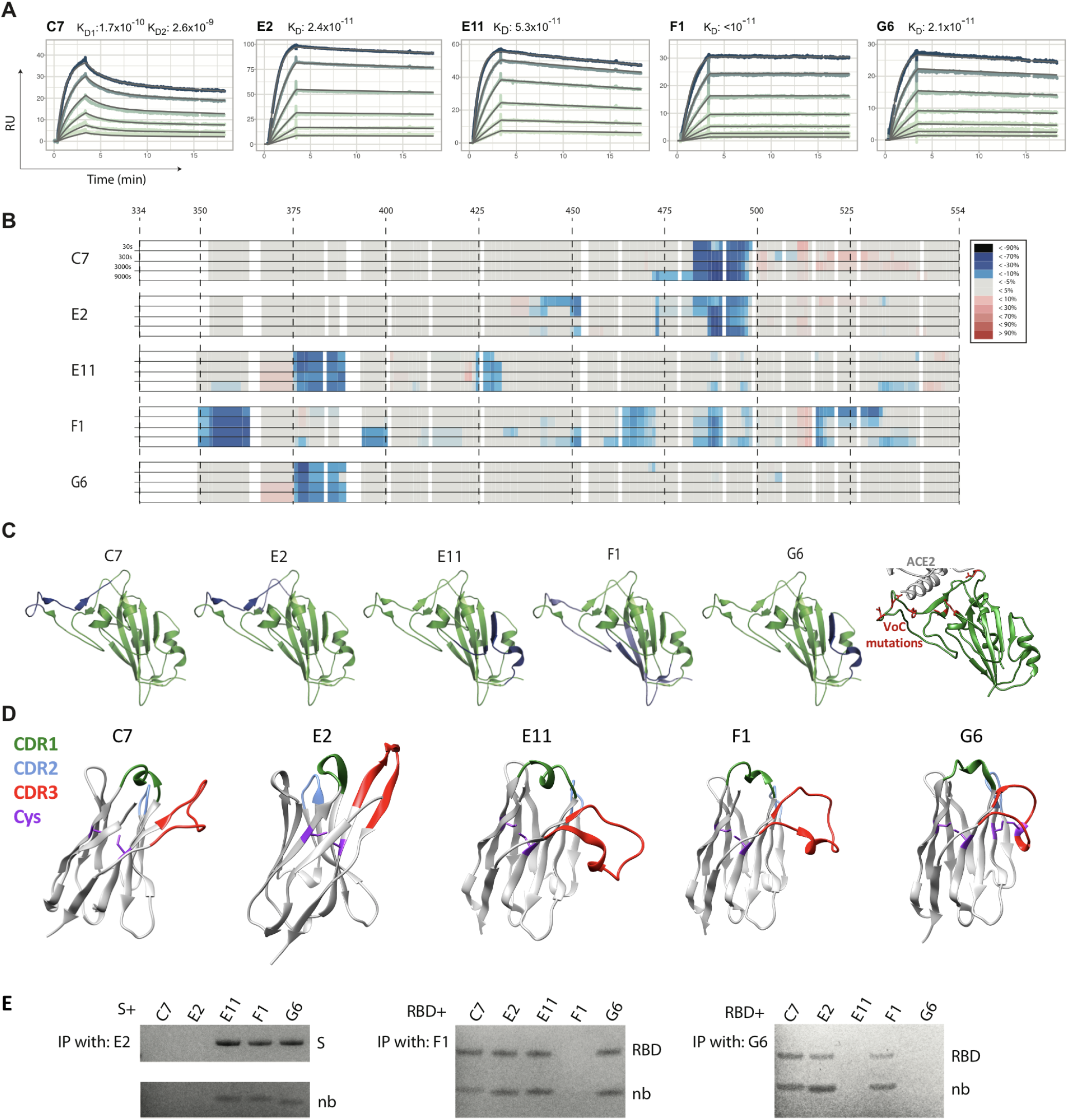
RBD-targeting affinities and epitope mapping. 5 RBD-targeting nanobodies were selected for affinity characterization by SPR and epitope mapping by HDX-MS. A shows picomolar affinities (170 pM to <10 pM) for all tested nanobodies. B depicts HDX-MS signal across the RBD sequence, which is mapped onto the RBD structure in C, together revealing three distinct epitope classes. Right-most in C shows the positions of all RBD mutations occurring in Variants Alpha, Beta, Gamma, Delta, Kappa, Epsilon, Eta, Iota and Lambda, shown in red. D shows AlphaFoldv2 ^25,26^ predictions of the structure for these five nanobodies, highlighting the CDRs, and the cysteine pairs. Unlike the others, G6 shows an additional predicted disulfide bond between cysteines in the CDR3 and in FR2. E shows immunoprecipitation competition analysis, supporting these three epitope classes. C7 and E2, the two most potent neutralizers of SARS-CoV-2 target an epitope at the ACE2 interface, which explains both their potency and their lack of cross reactivity. Nanobody G6 targets an epitope that is well conserved across the founder virus of SARS-CoV-2, the beta variant, and SARS-CoV-1 (where there is only a single substitution relative to SARS-CoV-2), explaining its cross reactivity, with E11 having a similar epitope. F1 has by far the largest epitope, which potentially explains the very low dissociation rate.

### RBD-specific nanobodies bind to different epitopes on the RBD

To determine the binding sites of some of the identified nanobodies, we used Hydrogen/Deuterium exchange coupled to mass spectrometry (HDX-MS). This technology provides a powerful means to study protein dynamics and interactions in solution ^27,28^. In the case of epitope mapping it can be viewed as a comparison of deuterium uptake between two states of a protein (unbound and bound), where the interaction leads to a change in conformational stability and/or solvent accessibility. In brief, nanobody-RBD complexes and RBD were exposed to deuterated H2O resulting in hydrogen/deuterium exchange, followed by denaturation and digestion with pepsin. Peptides were then analyzed by LC-MS. Comparison of bound and unbound RBD then allows to identify areas with altered HDX. All the investigated nanobodies were found to exhibit clear and defined interaction sites on the RBD (Fig. 5B,C). The structural resolution as defined by the degree of overlap of the peptides generated by pepsin digestion, i.e. the peptide map overlap, and the kinetics of the uptake is shown for respective state in supplementary file S1. Nanobodies E2 and C7 appear to have a common strong binding site spanning AA 487-496 where a strong protection was observed, with a unique weaker site for E2 spanning 441-452 and for C7 an extension of the strong binding site to include 471-486. F1 was the nanobody that introduced the most deuteration protection on the RBD, with a strong interaction at AA 352-361 but also at 392-400, 462-470, 483-492 and 514-530, explaining its exceptional affinity. When annotating these sites on a 3D structure of the RBD, it appears that F1 engages a large continuous surface. Finally, for G6 and E11 a common strong interaction site was observed spanning AA 375-387 and a unique E11 site at residues 423-431. Interestingly, G6, but not E11, has an additional cysteine pair connecting the CDR3 and FR2 (Fig. 5D). Although all nanobodies had at least one strong interaction site it should be noted that from the HDX data it is not possible to discriminate between a direct interaction surface and a possible conformational change in the RBD introduced by nanobody binding, when several interaction sites are observed. We confirmed the three distinct epitopes by immunoprecipitation-based competition assay (Fig. 5E).

Epitopes in the RBD have been broadly classified into 4 classes based on overlapping epitopes frequently targeted by antibodies isolated from convalescent humans ^29^. Nanobodies C7 and E2 were mapped to a class 2-like epitope, consistent with their ability to compete with ACE2 for binding to RBD, as well as the inability to neutralize the beta variant harbouring E484K. Nanobodies E11 and G6 were mapped to epitopes overlapping that of the monoclonal antibody COV2-2677^30,31^, consistent with that of a class 4 antibody (that also more broadly includes the antibody CR3022^32^). However, the ability of G6 but not E11 to prevent spike binding to ACE2 suggests it may utilize a different angle of approach. F1 appears to have a unique mode of recognition that does not map to any previously well-described epitopes.

### Therapeutic potential of nanobodies

To evaluate the therapeutic potential of neutralizing nanobodies for the treatment of SARS-CoV-2 infection, we used transgenic mice that express human ACE2 under the control of the cytokeratin-18 promoter (K18-hACE2 mice) ^33^. These mice are highly susceptible to SARS-CoV-2 infection, and experience weight loss following infection that correlates with pathology and disease severity ^34^. One major limitation to the therapeutic application of nanobodies is their short half-life in vivo. We therefore conjugated C7 to a nanobody specific for Albumin (Alb1) that has been demonstrated to increase serum halflife ^35^. Mice were challenged with 86 PFU of SARS-CoV-2 (2.4 × 10^6^ RNA genome copies) and subsequently treated with 320 μg C7-Alb1 (in 160 μl PBS) intraperitoneally (i.p.) on days 1 and 6 post infection. Untreated control mice experienced significant weight loss, beginning around 4 days following challenge. Both weight loss and viral load in oropharyngeal swabs were significantly lower for mice treated with C7-Alb1 compared to untreated mice (Fig. 6). Although three animals treated with C7-Alb1 experienced some transient weight loss between day 5 and 6, weight loss was significantly reduced compared to untreated mice (Fig. 6A,B). Furthermore, oropharyngeal viral loads in C7-Alb1 treated mice were approximately 100-fold lower than in untreated mice on day 5 (Fig. 6C). All but one of the untreated control mice succumbed to infection and had to be euthanized by day 7. However, all mice treated with C7-Alb1 survived, demonstrating the significant therapeutic efficacy of this neutralizing nanobody for SARS-CoV-2 infection.

**Fig. 6.**
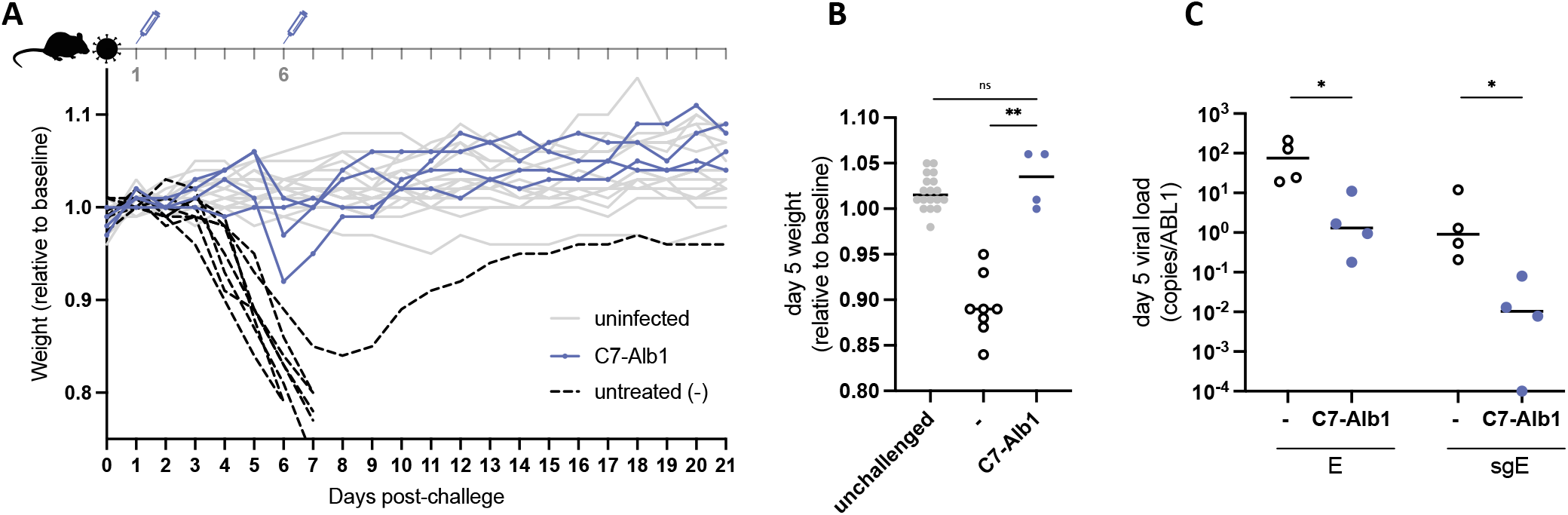
A half-life extended nanobody heterodimer rescues K18-hACE2 mice from a lethal SARS-CoV-2 challenge. K18-hACE2 mice were challenged with 86 PFU (2.4 *×* 10^6^ genome copies) of SARS-CoV-2, produced in Calu-3 cells, and weight was monitored over time (A). Four mice received 320 μg of C7-Alb1 i.p. at days 1 and 6 post-challenge. The mean weight of each mouse from day 0 to day 2 served as the baseline and the weight loss relative to this baseline is shown. Uninfected mice are shown in grey, untreated infected mice in black. B. Weight loss at day 5 post infection is shown for treated (C7-Alb1) and untreated (−) mice and compared to co-housed mice that were not challenged (unchallenged). ns, not significant (P>0.05); **, P<0.01. C. Viral load for both genomic (E) and subgenomic (sgE) RNA in oropharyngeal swabs taken at day 5 are shown for treated (C7-Alb1) and untreated (−) mice. *, P<0.05.

### Rapid click-chemistry based dimerization to identify potent homo- and heterodimer combinations

Nanobody dimerization can significantly increase potency ^36,37^. Combining nanobodies with distinct specificity can in addition limit viral escape ^8^. While molecular structures can be useful to select specific nanobody combinations, detailed structural information is not always readily available for larger pools of nanobodies. To identify potent dimers, we developed a screen to rapidly generate and test different nanobody combinations. We used sortase A functionalization and click chemistry to generate nanobody homo- and heterodimers as described in detail previously ^36^. In brief, using sortase A, nanobodies were C-terminally functionalized with a dibenzocyclooctyne (DBCO) or an azide. Labeled nanobodies were then mixed and incubated to permit sprain-promoted azide-alkyne cycloaddition (SPAAC) and C-to-C-terminal fusion of azide and DBCO-labeled nanobodies (Fig. 7A). Successful dimerization for all tested combinations was confirmed by SDS-PAGE analysis (Fig. S4), with dimerization efficiencies ranging between 55-70 %. Given that we were screening for substantial potency increases relative to the monomers, dimer reactions were not purified further before testing for their neutralization potential (Fig. 7B). The two most potent nanobodies C7 and E2 proved to be very potent in combination with any other nanobody. D9 and E11 proved to be good in combination with most other nanobodies. D4 and F12 displayed improved potency to varying degrees in combinations with the other nanobodies. F1 and G2, only performed well when combined with other potent neutralizers, and homo- or heterodimer combinations of these two appeared similar, or worse than the monomeric versions. For more precise concentration and neutralization value determination, homo- and heterodimers incorporating G6, E11 and E2 were produced and purified at larger scale (Fig. S2). All dimers proved extremely potent. An E2 homodimer with a solubility enhancing PEG11 linker neutralized SARS-CoV-2 with an IC50 of approx. 0.7 ng/ml (23 pM). E11 and G6 homo- and heterodimers neutralized with similar potency, between 10-20 ng/ml (333-667 pM). In conclusion, we show that high-throughput generation and screening of nanobody dimers can facilitate the rapid identification of extremely potent and synergistic combinations.

**Fig. 7.**
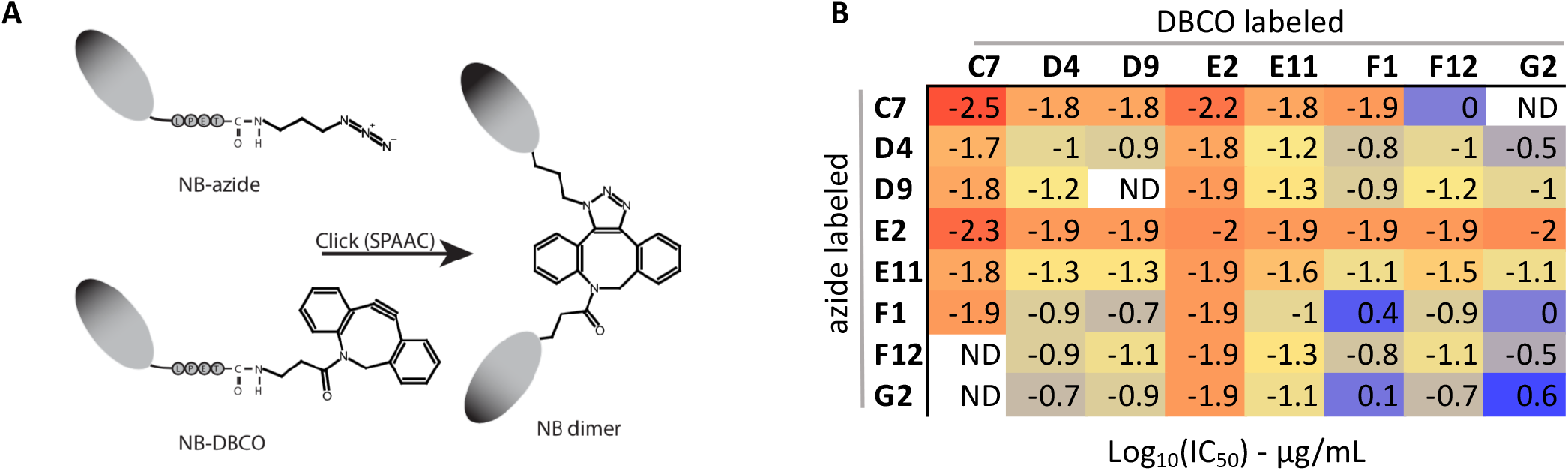
Rapid screening for potent nanobody dimer pairs. C-to-C-terminal fused nanobody dimers were generated using a combination of sortase A functionalization and click chemistry (A). B shows a heatmap of the neutralization IC50s against SARS-CoV-2 founder virus for crude homo- and heterodimer reaction products. Neutralization curves for select purified homo and heterodimers are shown in Fig. S2.

### Lineage-level analysis

Nanobodies do not exist as independent sequences, but as lineages arising from common VDJ recombination events that expand and undergo somatic hypermutation. Anti-spike single-domain antibodies from the same lineage are expected to bind the same epitope, but with varying affinities, and so identifying lineages can help guide both screening selection and downstream searches for optimized candidates. The grouping of sequence variants into lineage is not directly observed, and must be inferred from the sequences themselves. While the analysis of antibody lineages is growing in popularity in the antibody repertoire sequencing community, it has not been widely adopted for nanobodies, which may be, in part, due to the poor availability of germline sequence data. Many strategies for grouping lineages take all sequences with the same V and J germline assignments, and then cluster these based on the distance between their CDR3^38^. One of the benefits of relying on V and J assignments is reduced computational burden. The number of pairwise CDR3 comparisons is quadratic in the number of variants that must be compared, and splitting into groups by V and J assignment dramatically reduces the number of comparisons. Here we adopt a germline-naive lineage assignment strategy. Rather than V and J assignment, we use proximity in seqUMAP space to define a search neighbourhood over which to compare CDR3s (see Methods for a description of how to do this efficiently) and merge variants into lineages. Figure 8 explores the lineage structure of this nanobody dataset. In some cases, multiple nanobodies were screened from the same lineage. For example, panel D shows that three nanobodies, E11, G1, and F12, arise from distinct clades of a single lineage. These three nanobodies are up to 17 amino acids apart from each other, exhibit a 20-fold difference in potency between the most and least potent, and can cross neutralize the beta variant, and SARS-CoV-1. This lineage would be an excellent candidate for further screening - if another order of magnitude improvement were found in a variant on this lineage, this would be the most potent cross-neutralizing nanobody in the dataset.

**Fig. 8.**
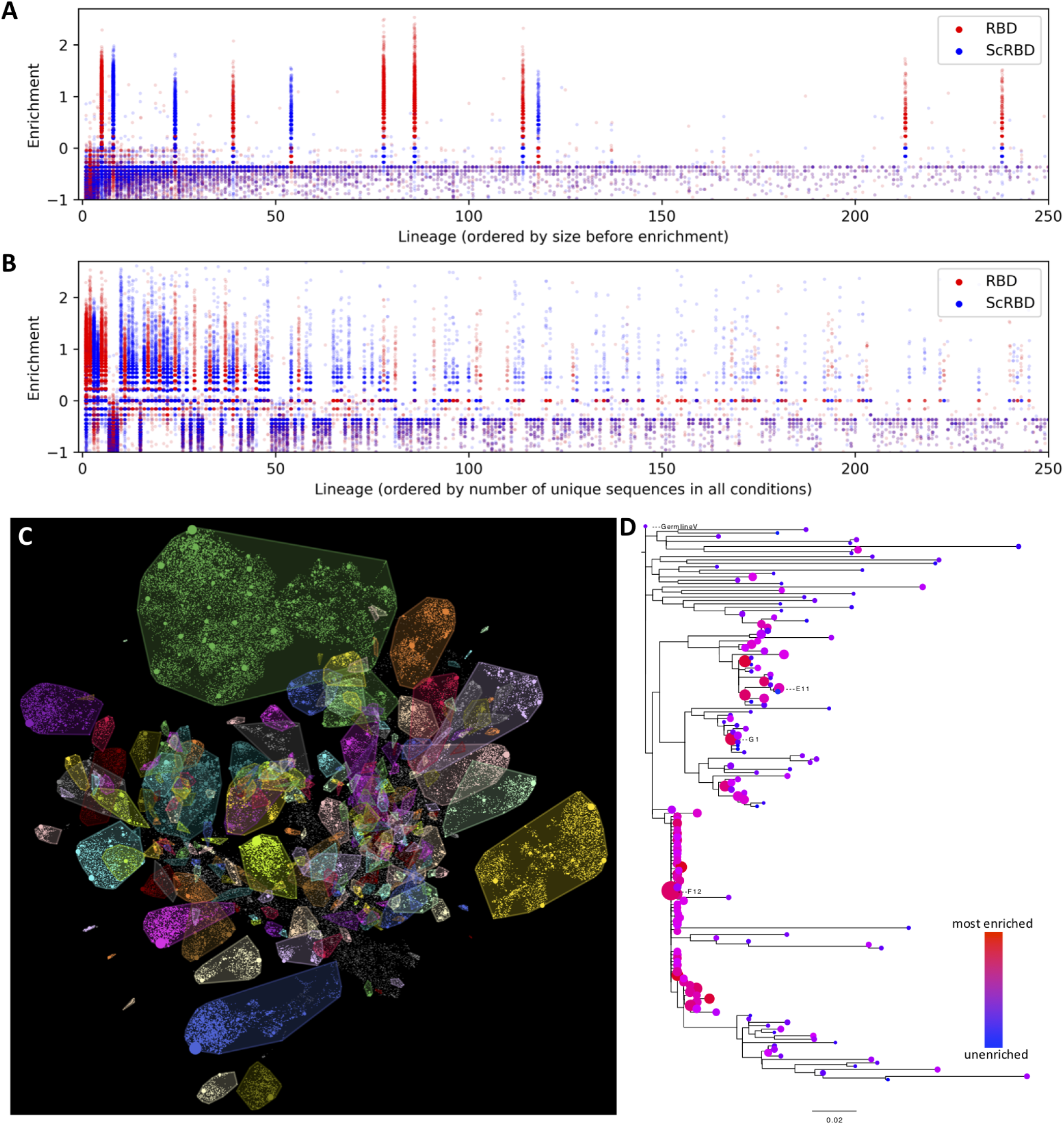
Nanobody lineages. Grouping nanobody variants by their inferred lineage, sharing a common original VDJ rearrangement, allows us to interrogate the clonality of the alpaca immune response, the consistency of enrichment, and lets us identify other candidate nanobody variants that might have improved properties. A and B shows the enrichment of variants, organized by lineage, in RBD and ScRBD enrichment conditions. Like at the variant level, enrichment at the lineage level is mutually exclusive, suggesting that lineages are each restricted to a single target. The lineage ordering in A shows that, prior to panning enrichment, only 11 of the largest 250 lineages are spike-specific, and B shows how this shifts with a single round of panning. C depicts lineages overlaid on the seqUMAP embeddings, and D shows the phylogeny for a subset the variants from a single lineage, containing nanobodies E11, G1, and F12. Neutralization data shows a 20-fold difference in potency between E11 and G1.

## Discussion

The use of nanobodies or nanobody-like proteins has increased greatly across a variety of applications in recent years. Although NGS-enabled analysis of immune repertoires is common for studying the elicitation and maturation of conventional antibodies ^38–42^, this approach is not yet standard for camelid repertoires, nor commonly used for nanobody discovery. Here, we describe a rapid and straight-forward approach to nanobody discovery that exploits the fact that, once established, nanobody libraries can be expanded indefinitely. This allows enrichment against multiple targets, which can be different sub-components or variants of the same antigen. Multiplexed NGS of the starting library and of each distinct enrichment step provides us with massively parallel information about the affinity of individual nanobodies against each target. In contrast to conventional panning and colony picking, this approach relies on enrichment metrics instead of post-panning abundance, which enables the identification of high affinity nanobodies even when they exist at low abundance in the baseline library, and are not sufficiently enriched to be sampled during traditional colony picking (see Fig. S5). Another benefit is that, as this approach relies on enrichment metrics, it only requires a single panning round. This could be expanded further to more complex antigens such as cell surfaces or different conformational states of proteins. Serendipitous discoveries have identified a number of nanobodies that recognize specific conformational states of proteins ^43,44^. By panning, for example, against different conformational states and identifying nanobodies that are enriched in one but not the other, our approach could provide the means to rationally isolate nanobodies with such conformation specificity.

This high-throughput approach provides joint enrichment and genetic information, and the genetic information allows grouping the nanobodies into lineages that will share functional properties. Most importantly, while their affinity may differ, nanobodies from the same lineage will target the same epitope. The advantages of a lineage-based analysis are two-fold: i) It avoids redundancy arising from testing multiple highly related nanobodies in the initial screen, and ii) once a promising lineage has been identified, information from across the lineage can be leveraged to identify or further optimize nanobodies with improvements in a particular function. We demonstrate the utility of the approach using a nanobody library obtained from an alpaca immunized with recombinant SARS-CoV-2 spike proteins, isolating a number of extremely potent, neutralizing nanobodies. These included nanobodies capable of neutralizing SARS-CoV-2 in the low picomolar range, as well as broadly neutralizing nanobodies capable of cross-neutralizing Variants of Concern and SARS-CoV-1.

Nanobodies E2 and C7 are among the most potent monomeric neutralizing nanobodies isolated to date, consistent with targeting of a ‘class 2’ epitope ^29^ overlapping the ACE2-binding site, which is highly represented among potently neutralizing antibodies isolated to date ^45,46^. Several neutralizing nanobodies have been evaluated against the beta variant, and the frequent resistance to neutralization suggests this is a frequent target for potently neutralizing nanobodies as well. Sziemel et al. ^47^ characterized three potent neutralizing nanobodies and, while one nanobody was similarly potent against beta, the other two neutralizing nanobodies failed to cross-neutralize this variant. Furthermore, a single mutation (E484K) was sufficient to recapitulate this escape. Similarly, in another nanobody library, the beta variant mutations (and E484K alone) were sufficient to almost completely abolish binding by two (out of 7) of the most potently neutralizing nanobodies ^48^.

F1, while not a potent neutralizer, shows exceptional binding properties that are of substantial benefit for diagnostic applications, where favorable binding may improve the ability to detect small quantities of SARS-CoV-2 antigen. The binding to a unique and conserved epitope, with associated cross-reactivity to the beta variant, is also desirable since diagnostic applications need to maintain sensitivity in a landscape dominated by such Variants of Concern.

G6, E11, and E4 are almost as potent as E2/C7, but exhibit robust cross-neutralization of the beta variant (B.1.351/501Y.V2), and no presently identified Variants of Concern harbour mutations in the epitopes identified for G6 and E11, suggesting that these may provide broad cross-variant neutralization. This property is becoming progressively more critical as Variants of Concern continue to emerge. For G6 and E11, this breadth is further supported by their neutralization of SARS-CoV-1, which has a single amino acid mutation in the epitope region common to G6 and E11, and one additional mutation in the E11-specific epitope region. Among sarbecoviruses that have the potential to use ACE2 (lacking two common deletions in the RBD that prevent ACE2 use ^49^), the overwhelming majority of these are identical in the G6 epitope to either SARS-CoV-1 or SARS-CoV-2, and since G6 neutralizes SARS-CoV-2 and tolerates the substitution in SARS-CoV-1, it is likely to exhibit substantial pan-sarbecovirus neutralization for ACE2-using sarbecoviruses.

We further show the therapeutic potential of nanobodies in a transgenic mouse model. A half-life extended C7 construct (C7-Alb1) administered following infection was able to rescue all treated animals from a near-universally fatal SARS-CoV-2 challenge dose. C7-Alb1 administered on day 1 and day 6 post-infection substantially reduced pathology, demonstrated by significantly reduced oropharyngeal viral loads and weight loss.

To be applied as therapeutics or prophylactics, nanobodies may need to undergo “humanization” to reduce their immunogenicity. One generally successful strategy is to replace the nanobody framework regions with those from related human immunoglobulin genes ^50^. With G6 in particular, the extreme variation in the CDR1 and CDR2 of enriched members of the G6 lineage suggest that binding may be completely dominated by the CDR3, suggesting humanization may be achievable without affecting neutralization potency. With its minimal binding footprint, a suitably humanized G6 construct would have exceptional potential as a broadly-neutralizing therapeutic or prophylactic against SARS-CoV-2 and its many emerging variants.

## ACKNOWLEDGEMENTS

We gratefully thank James Voss, Deli Huang, and Jesse Bloom for reagents. We thank Penny Moore and the NICD (South Africa) for providing the 501Y.V2 spike plasmid, which was generated using funding from the South African Medical Research Council. We thank Jonas Klingström for providing Calu-3 cells and sharing the infectious SARS-CoV-2 isolate. We thank Mateusz Kaduk for referring us to the AlphaFoldv2 implementation.

## METHODS

### Protein and probes

The plasmid for the expression of the SARS-CoV-2 prefusion-stabilized spike ^51^ was a kind gift from the McLellan lab. The plasmid was used for transient transfection of FreeStyle 293F cells using the FreeStyle MAX reagent (Thermo Fisher Scientific). The spike trimer was purified from filtered supernatant on Streptactin XT resin (IBA Lifesciences) or Ni-NTA resin and purified by size exclusion chromatography on a Superdex 200. The RBD domain was cloned upstream of a sortase A motif (LPETG) and a 6xHIS tag. The plasmid was used for transient transfection of FreeStyle 293F cells as described above. The protein was purified from filtered supernatant on His-Pur Ni-NTA resin followed by size-exclusion chromatography on a Superdex 200.

Nanobodies were cloned in the pHEN plasmid with a C-terminal sortase motif (LPETG) and a 6xHIS tag. BL21 E. coli were transformed with this plasmid, and expression was induced with 1 mM IPTG at OD600 = 0.6 and cells were grown overnight at 30 °C. Nanobodies were retrieved from the periplasm by osmotic shock and purified on Ni-NTA resin and size-exclusion chromatography.

The albumin binding nanobody Alb1 was described earlier ^35^ and the sequence was obtained from WO/2006/122787.

Sortase A 5M was produced as described before ^36^ in BL21 E. coli and purified by Ni-NTA and size exclusion chromatography.

Fluorescent spike ectodomain was generated by first attaching dibenzocyclooctyine-N-hydroxysuccinimidyl ester (DBCO-NHS) to the spike trimer in a 3:1 molar ratio, before attaching AbberiorStar-635P-azide by click-chemistry. The final product was purified from unreacted DBCO and fluorophore on a PD-10 desalting column. The biotinylated RBD was generated using sortase A and amine-PEG3-biotin as a nucleophile. The reaction was performed with 50 μM RBD, 5 μM sortase A 5M, and 8 mM amine-PEG3-biotin for 6 hours at 4 °C. Sortase A and unreacted RBD was removed on Ni-NTA resin and excess nucleophile was removed by two consecutive purifications on PD-10 desalting columns. The biotinylated nanobodies were generated using sortase with a reaction of 100 μM nanobody, 5 μM sortase A 5M, and 8 mM amine-PEG3-biotin for 2 hours at 25 °C. Sortase and unreacted nanobody was removed on Ni-NTA resin and the nanobodies were purified by size-exclusion chromatography or PD-10 desalting columns.

### Alpaca library generation and phage selection

The alpaca nanobody library used in this study was already described and used here ^19^. In brief, one adult female alpaca (Funny) at PreClinics, Germany, was immunized four times in a 60-day immunization schedule. Each immunization consisted of two injections. For the first immunization, 200 μg of prefusion stabilized spike and 200 μg of S1+S2 domain (Sino biologicals) was used. Remaining immunizations each consisted of one injection with 200 μg RBD and one injection with 200 μg prefusion stabilized spike, both produced in Freestyle 293F cells as described above. The animal study protocol was approved by the PreClinics animal welfare officer commissioner and registered under the registration No. 33.19-42502-05-17A210 at the Lower Saxony State Office for Consumer Protection and Food Safety—LAVES and is compliant with the Directive 2010/63/EU on animal welfare.

The nanobody phage library was generated as described by Hanke et al. ^6^. Phage display was performed on biotinylated RBD immobilized on streptavidin magnetic beads (Dynabeads M-280, Invitrogen) or strep-tagged spike immobilized on Strep-Tactin XT magnetic beads (IBA Lifesciences). To select for non-RBD binders, the phage enrichment on the spike protein was performed in the presence of non-immobilized RBD.

### Next generation sequencing and enrichment analysis

Nanobody phage libraries were sequenced at baseline and after each panning, as described in Hanke et al. ^6^. Briefly, phage libraries are amplified by PCR, and sequenced on an Illumina MiSeq instrument (2 × 300).

The establishment of a nanobody phage library described above overwrites the N-terminal region of the VHH with a cloning primer, which includes ambiguous nucleotides. Here we used this to investigate the internal consistency of each panning step. Sequencing primers were trimmed, and reads were collapsed by identity, retaining the frequency of each sequence. We performed an interim frequency analysis that compares the frequency of the two most frequent “versions” of each variant, where “versions” have identical sequences after the cloning primer, but distinct ambiguities in the cloning primer sequence (considering the three N-terminal ambiguities to prevent bias driven by priming effects). If a panning run is consistent in which variants are enriched, then there should be agreement between two versions of each variant. Figure S1 shows that this is true of RBD and ScRBD panning, but much less so of S panning. For downstream analysis, the cloning primer region was excluded when calculating variant frequencies. To reduce sequence error and the volume of data for downstream analysis, we exclude all non-functional sequences (with early stop codons), and all variants that do not occur at least three times across all datasets. Enrichment is estimated as the change in frequency between pre- and post-panning sequence datasets, in the log10 domain. To avoid undefined log ratios, we use pseudocount regularization when estimating frequencies. Enrichment is defined as: log_10_ (*F_post_*+ ε) - log_10_ (*F_pre_*+ ε), where *F_x_* is the proportion of reads from dataset *x*, and ε is a small constant, which we choose to be the reciprocal of the number of unique variants across the entire dataset.

Antibody repertoires exhibit complex relatedness patterns, due to the nature of VDJ recombination, and subsequent somatic hypermutation. To visualize the relatedness of sequences, we employ Uniform Manifold Approximation and Projection (UMAP ^17^). UMAP first constructs a neighbour graph in the original high-dimensional space, and then searches for a low-dimensional embedding that best preserves the similarity (measured by cross-entropy loss) between the original neighbour graph and the graph implied by the low-dimensional embedding. UMAP does not natively work on sequences (especially not unaligned sequences), and so we first embed sequences in a high dimensional “kmer” space, in which the squared euclidean distance between closely related sequences well approximates their levenshtein distance ^18^ especially for sequences that are closely related. As is popular in single-cell RNA seq applications of UMAP, we first project the high dimensional kmer representations into an intermediate space by PCA, and then apply UMAP to these PCA coordinates. The approach (which we call “seqUMAP”) is implemented in the Julia language and will be described in detail in a forthcoming manuscript. We apply seqUMAP to embed the set of 68,123 functional VDJ sequences into two dimensions.

### Lineages

Antibody repertoires are made up of distinct lineages. Within a lineage, antibodies share the same ancestor, but differ by somatic hypermutation. Clustering sequences into lineages typically relies on accurate V and J gene assignments, as the search for lineages is, for computational reasons, constrained to occur within sequences that share the same V and J germline gene. However, alpaca V gene databases appear to be incomplete, so we preferred an alternative strategy for lineage calling.

We begin with our seqUMAP embedding of all of the nanobody variants. We construct a “k-d tree” ^52^ of the seqUMAP coordinates of all variants, exploiting this space-partitioning data structure ^52^ to efficiently define a neighbour graph for all points within a set radius (0.4 here) of each other in seqUMAP space. The CDR3 is the strongest signal for lineage membership, and we consider each edge in G, and prune that edge if the CDR3s are too dissimilar. For efficient comparison of CDR3s, we again rely on kmer embeddings, and a “kmer distance” which approximates a length-normalized levenshtein distance ^18^. For CDR3s that are the same length, we prune edges where the kmer distance is greater than 12.5%, and for CDR3s of unequal length, we only allow up to 10% kmer distance. Most lineage calling strategies do not allow for any CDR3 length variation, which is partly for computational considerations, as this avoids a large number of pairwise alignment comparisons, but our kmer approach avoids any additional computational cost of comparing CDR3s of different lengths. Since CDR3 length variation is less common, and since allowing CDR3s of different lengths to be considered as the same lineage can lead to overly permissive lineage merging, we use a stricter distance requirement for CDR3s of unequal length. After CDR3-based pruning edges in G, the connected components of G define our single-domain antibody lineages.

This lineage calling relies on consistent CDR3 calling. We used an alignment based strategy to identify CDR3s. We constructed a multiple sequence alignment of all amino acid variants, and defined the CDR3 as the portion of the alignment after the canonical cysteine to the end of the conserved DYW, and project these regions back into nucleotide space, as the comparisons described above occur on CDR3 nucleotide sequences.

With the radii used for the analysis of our 68123 variant sequences, the exploitation of the seqUMAP neighbourhood reduces the number of pairwise CDR3 comparisons from 4.6 billion to 5.3 million, and the entire algorithm completes in around 15 seconds on a standard laptop.

### Cloning and expression of candidates

Selected nanobody sequences were ordered as eBlocks from Integrated DNA technologies (IDT) with 20 bp overhangs for Gibson assembly into a pHEN6 plasmid digested with PstI and BstEII restriction enzymes. The plasmids then encoded for nanobodies followed by a sortase A motif (LPETG) and a HIS tag. The Gibson assembly was performed in a 96-well plate and 2 μl of the assembly was directly used to transform BL21 E. coli and grown overnight in LB media and a 96-well plate covered with AirPore tape sheets in a 37 °C shaking incubator (>250 rpm). From this ‘master’ plate, 20 μl of culture was used to start an expression plate (1 ml LB per well in a 96-deep well plate). After 4 hours incubation at 37 °C, nanobody expression was induced by addition of 1 mM IPTG (final concentration), and cells were grown at 30 °C overnight, shaking. Cells in the expression plate were pelleted and resuspended in 100 μl TES buffer (200 mM Tris pH 8, 0.65 mM EDTA, 0.5 M sucrose) for 1 h. Resuspended cells were then diluted in 300 μl 0.25 × TES buffer overnight. Cells were centrifuged, periplasmic extracts collected, and directly used for expression level quantification, ELISA and neutralization assays.

For nanobody candidates that were analysed in more detail, plasmids from the original Gibson assembly were amplified in DH5α and verified by sanger sequencing, before being produced in larger quantities and purified by Ni-NTA affinity and size-exclusion chromatography.

### Neutralization assay

Pseudoviruses were generated by co-transfection of HEK293T cells with plasmids encoding firefly luciferase, a lentiviral packaging plasmid (Addgene cat8455), and a plasmid encoding the spike protein (with a C-terminal truncation) from either SARS-CoV (Addgene cat 170447), SARS-CoV-2 ^53^, or SARS-CoV-2 B.1.351 variant (beta) ^23^. Media was changed 12-16 h post-transfection, and pseudotyped viruses were harvested at 48- and 72 hours, clarified by centrifugation and stored at −80 °C until use. Pseudotyped viruses (PSV) sufficient to generate 100 000 relative light units (RLU) were incubated with serial dilutions of nanobody for 60 min at 37 °C. 15 000 HEK293T-hACE2 cells were then added to each well, and the plates were incubated for 48 h at 37 °C. Luminescence was measured using Bright-Glo (Promega) on a GM-2000 luminometer (Promega) with an integration time of 0.3 s. Neutralizing antibody ID50 titers were calculated in Prism 9 (GraphPad Software) by fitting a four-parameter logistic curve bounded between 0 and 100, and interpolating the concentration/dilution where RLUs were reduced by 50% relative to control wells in the absence of nanobody.

For periplasmic extract PSV neutralization, low-level background inhibition was evident. For the rapid screen, we subtracted this background value from all IC50 for the results described in Fig. 3, and for candidate selection.

### Flow cytometry

HEK293T-hACE2 cells were trypsinized and fixed in 4% formaldehyde in PBS for 20 min. Cells were stained with spike-AbberiorStar635P not premixed or premixed with target nanobody or control nanobody. Fluorescence was quantified using a BD FACSCelesta and the FlowJo software package.

### Surface plasmon resonance

Binding kinetics were determined by surface plasmon resonance using a BIAcore 2000. All experiments were performed at 25 °C in a running buffer of 10 mM HEPES, 150 mM NaCl, pH 7.4, and 0.005% Tween-20 (v/v). Site-specifically biotinylated RBD was immobilized on streptavidin sensor chips (Series S sensor Chip SA, GE Healthcare) to a level of 200 resonance units (RU). A 2-fold dilution series of the nanobodies was injected at a flow rate of 30 μl/min (association 180 s, dissociation 900 s), and the immobilized RBD was regenerated using 0.1 M glycine buffer pH 2 for 2x 10 seconds. Data were analyzed using BIAevaluation Software and fitted using the 1:1 Langmuir model with mass transfer, except for C7 where we used the heterogeneous ligand model to account for the self-dimerization of this nanobody.

### Epitope mapping by Hydrogen–deuterium exchange mass spectrometry

All chemicals were from Sigma Aldrich, pH measurements were made using a SevenCompact pH-meter equipped with an InLab Micro electrode (Mettler-Toledo), prior to all measurements a 4 point calibration (pH 2,4,7,10) was made. The HDX-MS analysis was made using automated sample preparation on a LEAP H/D-X PAL™ platform interfaced to an LC-MS system, comprising an Ultimate 3000 micro-LC coupled to an Orbitrap Q Exactive Plus MS.

HDX was performed on an RDB, 0.6 mg/ml without and with nanobody (E2, C7, F1, G6 and E11), in PBS, pH 7.5 (Sigma-Aldrich, D8537). All HDX-MS was done in one continuous run, with runs of the apo state made in between the nanobody runs, in total 8 replicates were made for the apo state, for E2, C7, F1, E11 triplicates and for EB-G6 duplicate samples were run. Apo state samples constituted 2.5 μl RDB and 2.5 μl PBS and the interaction analysis samples, 2.5 μl RDB mixed with 2.5 μl ligand, the samples were diluted with 30 μl 10 mM PBS, pH 7,46 or HDX labelling buffer of the same composition prepared in D2O, pH(read) 7.10. The HDX labelling was carried out for t = 0, 30, 300, 3000 and 9000s at 20°C. The labelling reaction was quenched by dilution of 30 μl labelled sample with 30 μl of 1% TFA, M TCEP, 4 M urea, pH 2.5 at 1°C, 60 μl of the quenched sample was directly injected and subjected to online pepsin digestion at 4 °C (in-house immobilized pepsin column, 2.1 × 30 mm). The online digestion and trapping was performed for 4 minutes using a flow of 50 μL/min 0.1 % formic acid, pH 2.5. The peptides generated by pepsin digestion were subjected to on-line SPE on a PepMap300 C18 trap column (1 mm × 15 mm) and washed with 0.1% FA for 60s. Thereafter, the trap column was switched in-line with a reversed-phase analytical column, Hypersil GOLD, particle size 1.9 μm, 1 × 50 mm, and separation was performed at 1°C using a gradient of 5-50 % B over 8 minutes and then from 50 to 90% B for 5 minutes, the mobile phases were 0.1 % formic acid (A) and 95 % acetonitrile/0.1 % formic acid (B). Following the separation, the trap and column were equilibrated at 5% organic content, until the next injection. The needle port and sample loop were cleaned three times after each injection with mobile phase 5%MeOH/0.1%FA, followed by 90% MeOH/0.1%FA and a final wash of 5%MeOH/0.1%FA. After each sample and blank injection, the Pepsin column washed by injecting 90 μl of pepsin wash solution 1% FA /4 M urea /5% MeOH. In order to minimize carry-over a full blank was run between each sample injection. Separated peptides were analysed on a Q Exactive Plus MS, equipped with a HESI source operated at a capillary temperature of 250 °C with sheath gas 12, Aux gas 2 and sweep gas 1 (au). For HDX analysis MS full scan spectra were acquired at 70K resolution, AGC 3e6, Max IT 200ms and scan range 300-2000. For identification of generated peptides separate undeuterated samples were analysed using data dependent MS/MS with HCD fragmentation.

A summary of the HDX experimental detail is reported in Supplementary File 1. The mass spectrometry and HDExaminer analysis files have been deposited to the ProteomeXchange Consortium via the PRIDE partner repository (REF ID:30395289).

### HDX-MS Data analysis

PEAKS Studio X Bioinformatics Solutions Inc. (BSI, Waterloo, Canada) was used for peptide identification after pepsin digestion of undeuterated samples. The search was done on a FASTA file with only the RDB sequence, search criteria was a mass error tolerance of 15 ppm and a fragment mass error tolerance of 0.05 Da, allowing for fully unspecific cleavage by pepsin. Peptides identified by PEAKS with a peptide score value of log P > 25 and no modifications were used to generate a peptide list containing peptide sequence, charge state and retention time for the HDX analysis. HDX data analysis and visualization was performed using HDExaminer, version 3.1.1 (Sierra Analytics Inc., Modesto, US). Each nanobody + RDB state was analysed and compared to its four closest (in time) apo state runs. The analysis was made on charge states 1-6 for each peptide, allowed only for EX2 and the two first residues of a peptide were assumed unable to hold deuteration. Due to the comparative nature of the measurements, the deuterium incorporation levels for the peptic peptides were derived from the observed relative mass difference between the deuterated and non-deuterated peptides without back-exchange correction using a fully deuterated sample ^54^. As a full deuteration experiment was not made, full deuteration was set to 75% of maximum theoretical uptake. The presented deuteration data is the average of all high and medium confidence results. The allowed retention time window was ± 30 seconds. Heatmaps settings were uncoloured proline, heavy smoothing and the difference heatmaps were drawn using automatically calculated significance based on replicate variance. The spectra for all time points were manually inspected; low scoring peptides, obvious outliers and any peptides where retention time correction could not be made consistent were removed. As bottom-up labelling HDX-MS is limited in structural resolution by the degree of overlap of the peptides generated by pepsin digestion, the peptide map overlap is shown for respective state in Supplementary file 1.

### Nanobody competition assay

Immunoprecipitations for competition assays were performed with 8 μg of C-terminally biotinylated nanobody on M-280 streptavidin magnetic beads (Dynabeads, Invitrogen) and 10 μg of spike or RBD preincubated with 10 μg of the indicated (HIS-tagged) nanobodies. Bound spike or RBD was eluted in 0.2 M glycine pH 2.2 and analyzed by SDS-PAGE and Coomassie staining.

### SARS-CoV-2 challenge experiments

K18-hACE2 transgenic mice were purchased from Jackson laboratories and maintained as a hemizygous line. Experiments were conducted in BSL3 facilities at the Comparative Medicine department (KM-F) at Karolinska Institutet. Ethics for studies of virus infection and therapeutic intervention were obtained from the Swedish Board of Agriculture (10513-2020). Mice were administered nanobodies as described in the main text and challenged intranasally with 86 PFU SARS-CoV-2 in 40 μl PBS following isoflurane sedation. Oropharyngeal sampling was performed on day 5, under light anesthesia with isoflurane. Weight and general body condition were monitored daily until weight drop started, whereupon mice were monitored twice daily. During the experiment, weight loss, changes in general health, breathing, body movement and posture, piloerection and eye health were monitored. Mice were sacrificed when they reached 20% weight loss or when movement was greatly impaired and/or they experienced difficulty breathing that was considered to reach a severity level of 0.5 on Karolinska Institutet’s veterinary plan for monitoring animal health. The weight loss in response to infection was highly reproducible. In Fig. 6 data from 50% of the untreated, challenged animals are historical controls from previous experiments performed under identical conditions. This challenge experiment was run at the same time as that from Hanke et al. (2021) ^19^, and the control mice were shared between both.

### RNA Extraction and RT-qPCR for SARS-CoV-2 detection

Viral RNA was isolated from buccal swabs collected 5 days post infection and stored in 500 μl of TRIzol™ Reagent (Invitrogen). Total RNA extractions from buccal swab samples were performed using an adapted TRIzol™ manufacturers protocol with a 45 min precipitation step at −20° C. RNA pellets were resuspended in 20 μl of RNase-free water. RT-PCR reactions were performed using 4 μl of resuspended RNA in a 20 μl reaction volume using the Superscript III one step RT-qPCR system with Platinum Taq Polymerase (Invitrogen) with 400 nM concentrations of each primer and 200 nM of probe. Primers and probes for the CoV-E gene target were as previously described ^55^. Primers and probes for the ABL1 target were adapted from Ishige et al. ^56^ to enable detection of the murine homolog: ABL1-ENF1003-deg: 5’-TGGAGATAACACTCTCAGCATKACTAAAGGT-3’ ABL1-ENR1063: 5’-GATGTAGTTGCTTGGGACCCA-3’ ABL1-ENPr1043-deg: 5’-HEX-CCATTTTTSGTTTGGGCTTCACACCATT-BHQ1-3’. The CoV-E and ABL1 TaqMan assays were run in multiplex. Detection of the subgenomic CoV-E target was adapted from Wölfel et al. ^57^, using a leader/E gene junction specific forward primer: sgEjunc-SARSCoV2-F 5’-CGATCTCTTGTAGATCTGTTCTCTAAACG-3’ All oligonucleotides were synthesized by Eurofins Genomics.

Thermal cycling conditions for all assays consisted of RT at 55 °C for 10 min, denaturation at 95 °C for 3 min, and 45 cycles of 95 °C, 15s and 58 °C, 30s. Reactions were carried out using a CFX96 Connect Real-Time PCR Detection System (Bio-Rad) following manufacturer instructions. To generate standard curves, a synthetic DNA template gBlock (Integrated DNA Technologies) was transcribed using the mMessage mMachine™ T7 Transcription Kit (Invitrogen) and serially diluted. To reduce sampling-related variability, SARS-CoV-2 RNA copies were normalized by ABL1 copies, and this ratio was used for comparisons. ABL1 copies were not significantly different between groups.

### Generation of nanobody dimers by sortase-mediated functionalization and click chemistry

Nanobodies were functionalized site-specifically on the C-terminus using sortase A 5M with either an azide or a dibenzocyclooctyne (DBCO) and subsequently dimerized by Cu-free strain-promoted azide-alkyne click chemistry (SPAAC) reaction as described earlier ^36^. In brief, nanobodies at concentrations ranging from 75 μM to 205 μM were incubated with 5 μM sortase A, 8 mM DBCO-amine (Sigma–Aldrich, 761540) or 10 mM 3-Azido-1-propanamine (Sigma–Aldrich, 762016), in 50 mM Tris pH 7.5, 150 mM NaCl, 10 mM CaCl2, for 3 h at 25 °C. Unreacted nanobody, sortase A and excess nucleophile were removed using Ni-NTA resin and Zeba spin desalting columns (2 mL, 7K MWCO, Thermo Fisher Scientific, 89890). Dimers were generated with a SPAAC reaction by incubating 30 μg of DBCO functionalized and 30 μg of azide functionalized nanobody in a 96-well plate for 72 hours at 4 °C. Reaction products were analyzed by SDS-PAGE (4–12% NuPAGE Bis-Tris, Life Technologies) and Coomassie G-250 staining. The relative amount of dimers in the gels (Fig. S4) was quantified using ImageJ and an E4 homodimer as a reference. Some homodimers, when produced at larger scale (Fig. S2), were generated by combining azide-functionalized nanobody with bis-PEG11-DBCO to increase solubility. These constructs were purified by size-exclusion chromatography.

---

**Fig. SI1.**
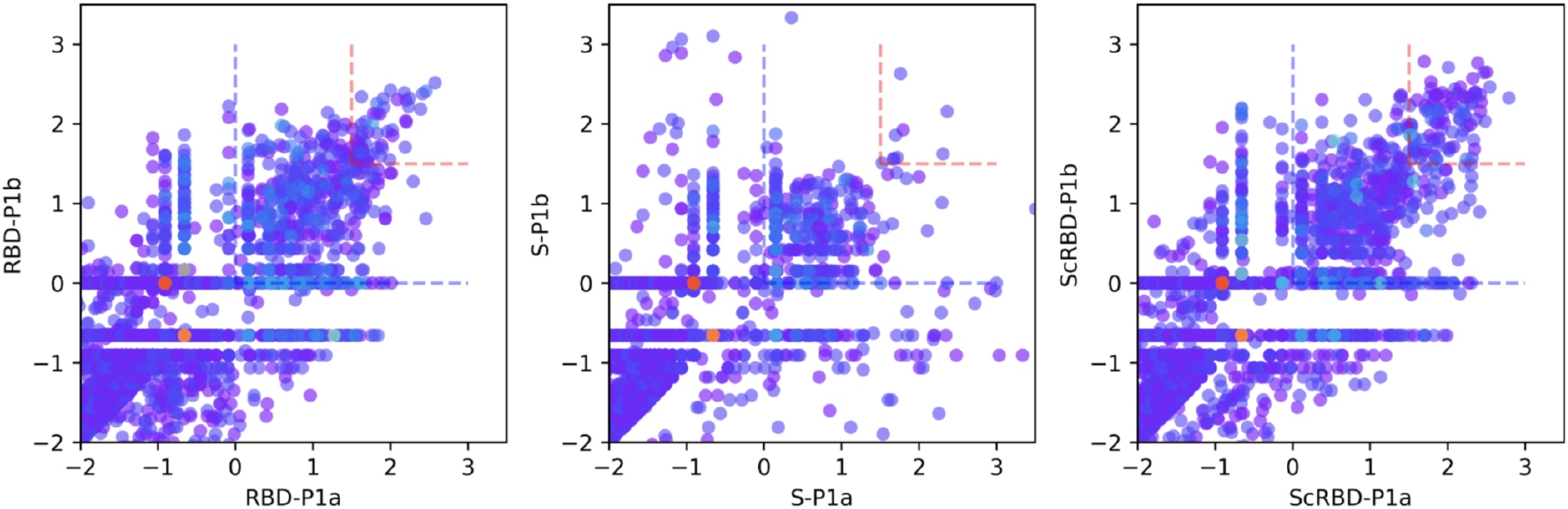
Within-panning consistency estimation. We compare two “versions” of each nanobody variant that differ by degenerate bases introduced with the cloning primer when establishing the baseline library. RBD and ScRBD panning shows consistency in enrichment, but spike panning is less consistent.

**Fig. SI2.**
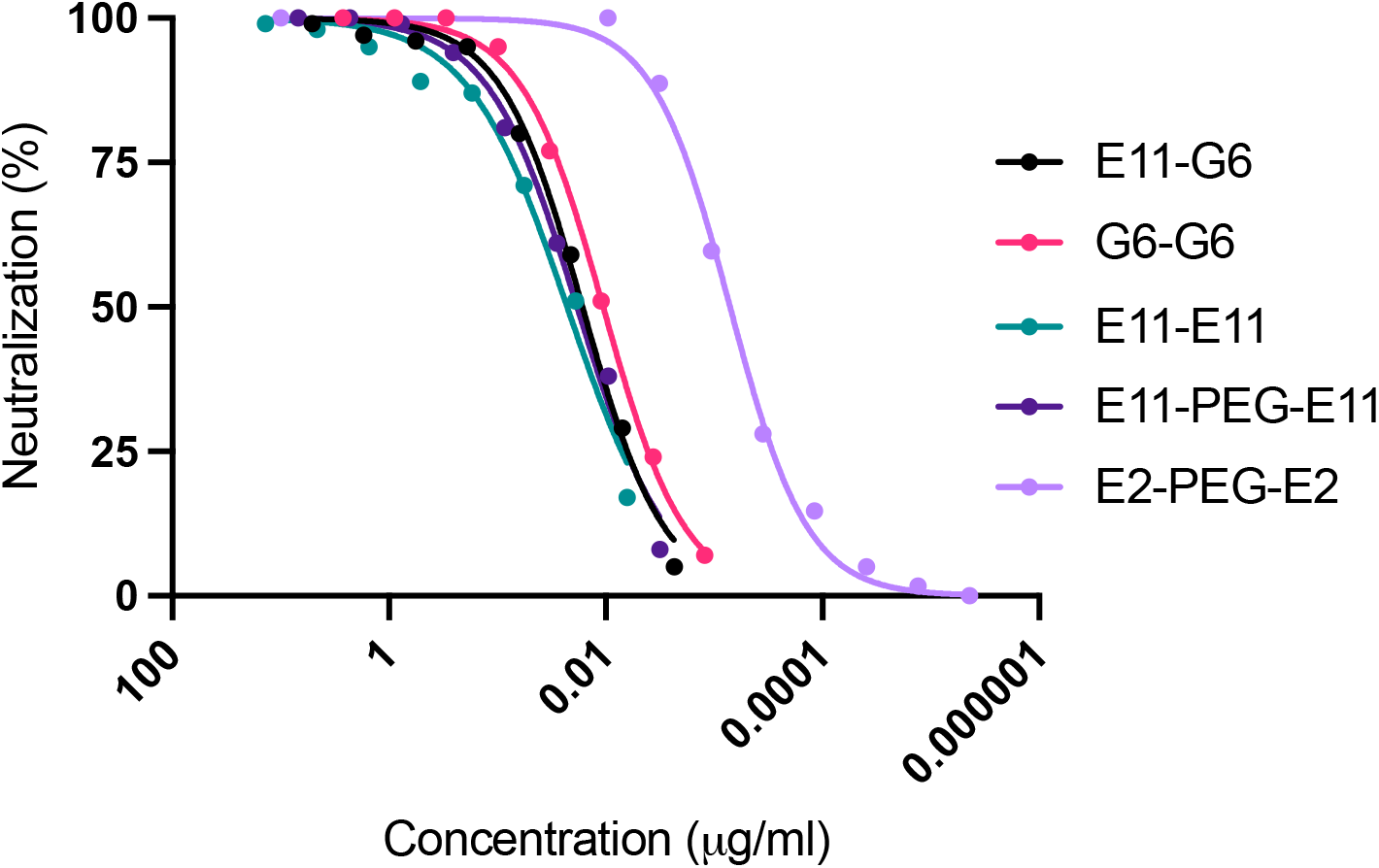
Ultrapotent neutralization of SARS-CoV-2 by nanobody homo- and heterodimers. Neutralization of SARS-CoV-2 spike pseudotyped lentiviruses are shown for nanobody dimers generated by Cu-free strain-promoted azide-alkyne click chemistry (SPAAC) reaction. E11 and G6 homo- and heterodimers neutralized with similarly high potency. An E2 dimer (with a short PEG11 spacer, E2-PEG-E2) was ultrapotent, with an IC50 of approximately 0.7 ng/ml (23 pM).

**Fig. SI3.**
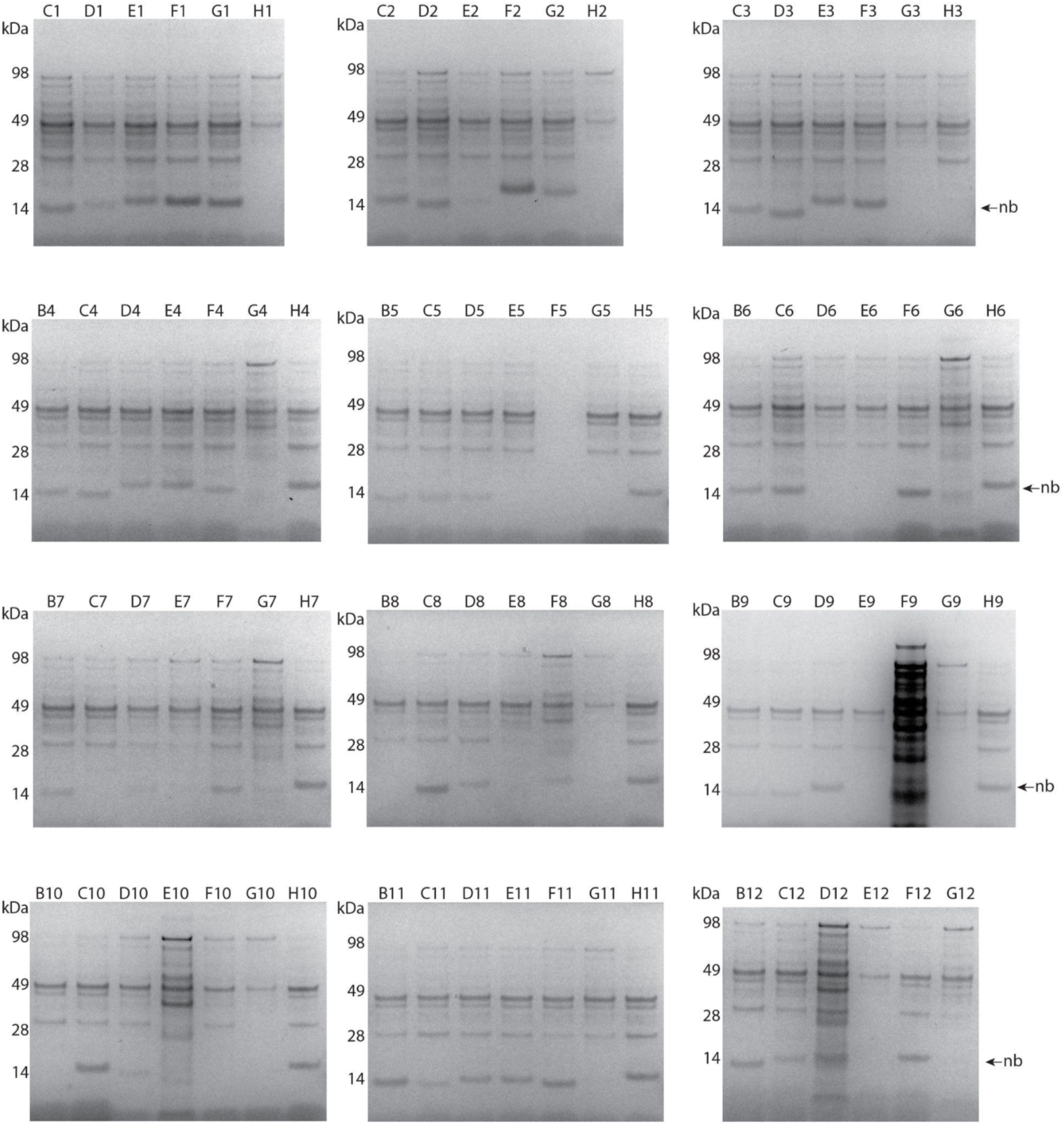
SDS-PAGE analysis and Coomassie staining of periplasmic extracts. Nanobodies (nb) run at 15 kDa.

**Fig. SI4.**
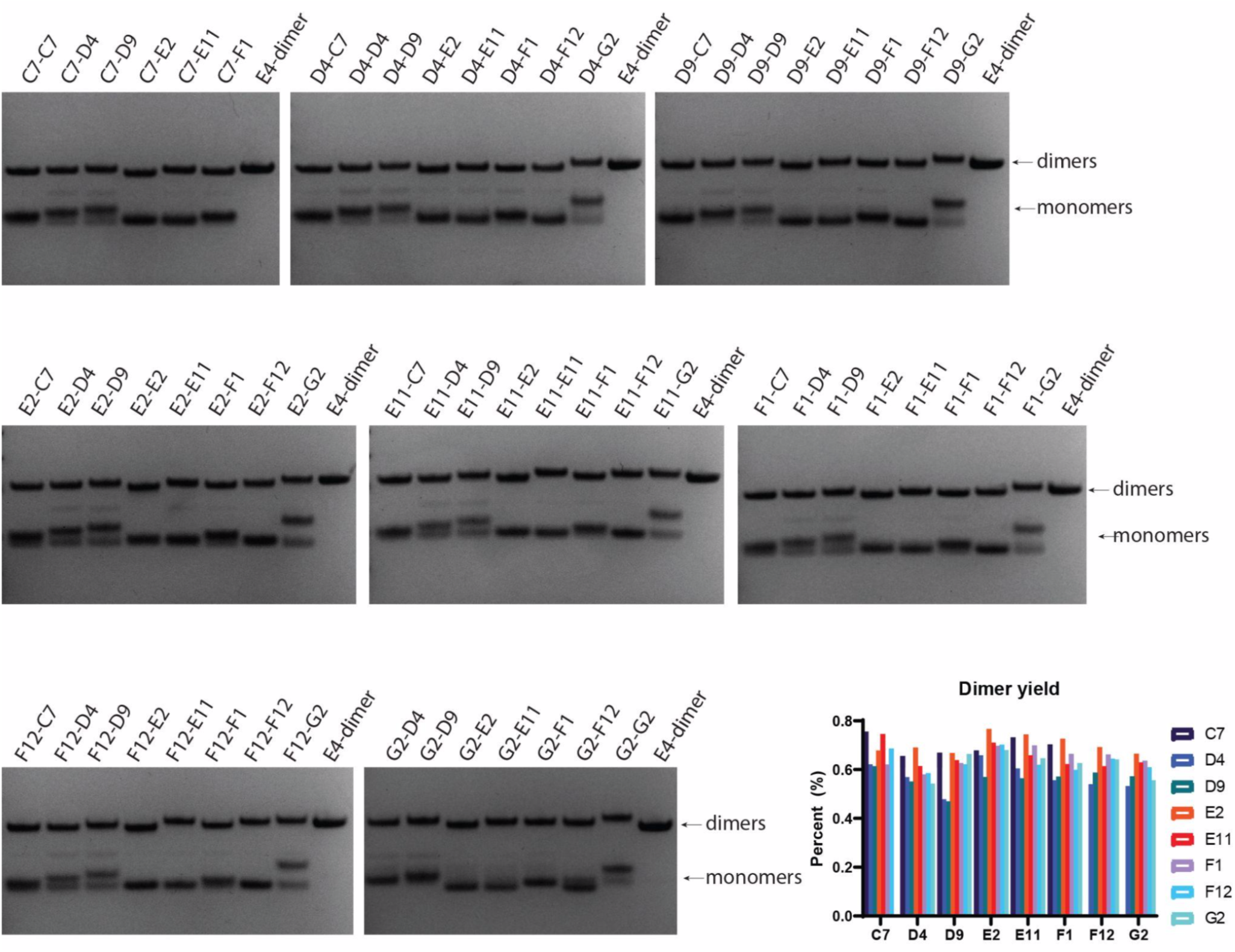
SDS-PAGE analysis and Coomassie staining of nanobody dimers generated to rapidly screen for potently neutralizing dimer pairs.

**Fig. SI5.**
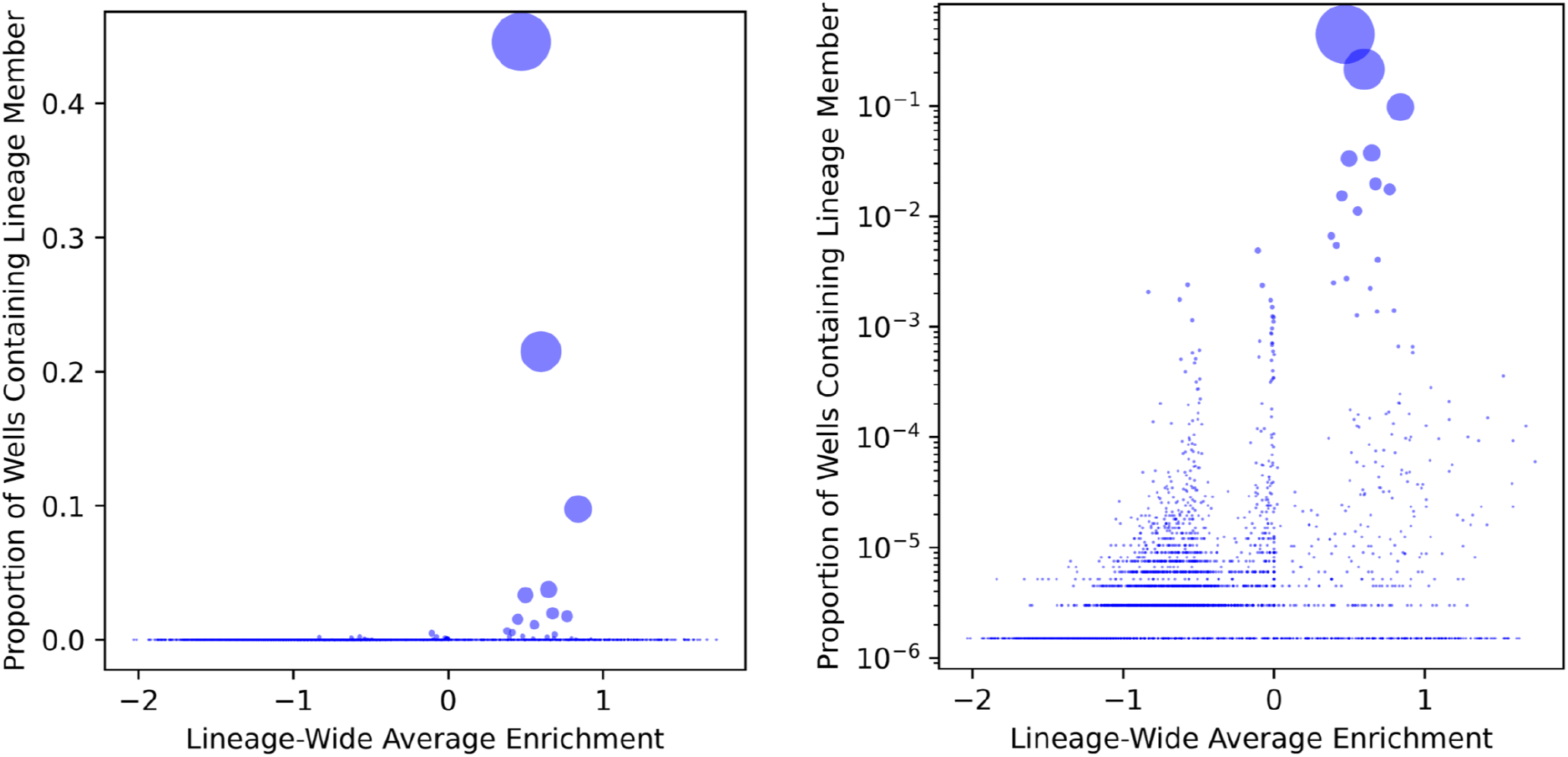
Lineage enrichment vs sampling probability. Each bubble represents a lineage, showing the mean enrichment for each lineage vs the probability that a single well would contain a member from that lineage, after a second round of RBD panning, estimated from NGS frequency data (left: linear scale; right: log scale). The skewed size distribution of lineages, where a small number of lineages are extremely high frequency, shows that colony-based sampling approaches will not reach the lower frequency lineages. As an example, we would have had to screen nearly 500 colonies to have a >50% chance of discovering a member from the lineage of our highest affinity RBD binding nanobody (F1).

## Bibliography

1. C Hamers-Casterman, T Atarhouch, S Muyldermans, G Robinson, C Hamers, E B Songa, N Bendahman, and R Hamers. Naturally occurring antibodies devoid of light chains. Nature, 363(6428):446–448, June 1993.

2. Sarah A Nordeen, Kasper R Andersen, Kevin E Knockenhauer, Jessica R Ingram, Hidde L Ploegh, and Thomas U Schwartz. A nanobody suite for yeast scaffold nucleoporins provides details of the nuclear pore complex structure. Nat. Commun., 11(1):6179, December 2020.

3. Tomasz Uchański, Simonas Masiulis, Baptiste Fischer, Valentina Kalichuk, Uriel López-Sánchez, Eleftherios Zarkadas, Miriam Weckener, Andrija Sente, Philip Ward, Alexandre Wohlkönig, Thomas Zögg, Han Remaut, James H Naismith, Hugues Nury, Wim Vranken, A Radu Aricescu, Els Pardon, and Jan Steyaert. Megabodies expand the nanobody toolkit for protein structure determination by single-particle cryo-EM. Nat. Methods, 18(1):60–68, January 2021.

4. Yushu Joy Xie, Michael Dougan, Noor Jailkhani, Jessica Ingram, Tao Fang, Laura Kummer, Noor Momin, Novalia Pishesha, Steffen Rickelt, Richard O Hynes, and Hidde Ploegh. Nanobody-based CAR T cells that target the tumor microenvironment inhibit the growth of solid tumors in immunocompetent mice. Proc. Natl. Acad. Sci. U. S. A., 116(16):7624–7631, April 2019.

5. Novalia Pishesha, Thibault Harmand, Liyan Y Smeding, Weiyi Ma, Leif S Ludwig, Robine Janssen, Ashraful Islam, Yushu J Xie, Tao Fang, Nicholas McCaul, William Pinney, 3rd, Harun R Sugito, Martin A Rossotti, Gualberto Gonzalez-Sapienza, and Hidde L Ploegh. Induction of antigen-specific tolerance by nanobody-antigen adducts that target class-II major histocompatibility complexes. Nat Biomed Eng, June 2021.

6. Leo Hanke, Laura Vidakovics Perez, Daniel J Sheward, Hrishikesh Das, Tim Schulte, Ainhoa Moliner-Morro, Martin Corcoran, Adnane Achour, Gunilla B Karlsson Hedestam, B Martin Hällberg, Ben Murrell, and Gerald M McInerney. An alpaca nanobody neutralizes SARS-CoV-2 by blocking receptor interaction. Nat. Commun., 11(1):4420, September 2020.

7. Jiandong Huo, Audrey Le Bas, Reinis R Ruza, Helen M E Duyvesteyn, Halina Mikolajek, Tomas Malinauskas, Tiong Kit Tan, Pramila Rijal, Maud Dumoux, Philip N Ward, Jingshan Ren, Daming Zhou, Peter J Harrison, Miriam Weckener, Daniel K Clare, Vinod K Vogirala, Julika Radecke, Lucile Moynié, Yuguang Zhao, Javier Gilbert-Jaramillo, Michael L Knight, Julia A Tree, Karen R Buttigieg, Naomi Coombes, Michael J Elmore, Miles W Carroll, Loic Carrique, Pranav N M Shah, William James, Alain R Townsend, David I Stuart, Raymond J Owens, and James H Naismith. Neutralizing nanobodies bind SARS-CoV-2 spike RBD and block interaction with ACE2. Nat. Struct. Mol. Biol., 27(9):846–854, September 2020.

8. Paul-Albert Koenig, Hrishikesh Das, Hejun Liu, Beate M Kümmerer, Florian N Gohr, Lea-Marie Jenster, Lisa D J Schiffelers, Yonas M Tesfamariam, Miki Uchima, Jennifer D Wuerth, Karl Gatterdam, Natalia Ruetalo, Maria H Christensen, Caroline I Fandrey, Sabine Normann, Jan M P Tödtmann, Steffen Pritzl, Leo Hanke, Jannik Boos, Meng Yuan, Xueyong Zhu, Jonathan L Schmid-Burgk, Hiroki Kato, Michael Schindler, Ian A Wilson, Matthias Geyer, Kerstin U Ludwig, B Martin Hällberg, Nicholas C Wu, and Florian I Schmidt. Structure-guided multivalent nanobodies block SARS-CoV-2 infection and suppress mutational escape. Science, 371(6530), February 2021.

9. Bert Schepens, Loes van Schie, Wim Nerinckx, Kenny Roose, Wander Van Breedam, Daria Fijalkowska, Simon Devos, Wannes Weyts, Sieglinde De Cae, Sandrine Vanmarcke, Chiara Lonigro, Hannah Eeckhaut, Dries Van Herpe, Jimmy Borloo, Ana Filipa Oliveira, Joao Paulo Catani, Sarah Creytens, Dorien De Vlieger, Gitte Michielsen, Jackeline Cecilia Zavala Marchan, George D Moschonas, Iebe Rossey, Koen Sedeyn, Annelies Van Hecke, Xin Zhang, Lana Langendries, Sofie Jacobs, Sebastiaan ter Horst, Laura Seldeslachts, Laurens Liesenborghs, Robbert Boudewijns, Hendrik Jan Thibaut, Kai Dallmeier, Greetje Vande Velde, Birgit Weynand, Julius Beer, Daniel Schnepf, Annette Ohnemus, Isabel Remory, Caroline S Foo, Rana Abdelnabi, Piet Maes, Suzanne J F Kaptein, Joana Rocha-Pereira, Dirk Jochmans, Leen Delang, Frank Peelman, Peter Staeheli, Martin Schwemmle, Nick Devoogdt, Dominique Tersago, Massimiliano Germani, James Heads, Alistair Henry, Andrew Popplewell, Mark Ellis, Kevin Brady, Alison Turner, Bruno Dombrecht, Catelijne Stortelers, Johan Neyts, Nico Callewaert, and Xavier Saelens. Drug development of an affinity enhanced, broadly neutralizing heavy chain-only antibody that restricts SARS-CoV-2 in rodents. March 2021.

10. David R Maass, Jorge Sepulveda, Anton Pernthaner, and Charles B Shoemaker. Alpaca (lama pacos) as a convenient source of recombinant camelid heavy chain antibodies (VHHs). J. Immunol. Methods, 324(1-2):13–25, July 2007.

11. Els Pardon, Toon Laeremans, Sarah Triest, Søren G F Rasmussen, Alexandre Wohlkönig, Armin Ruf, Serge Muyldermans, Wim G J Hol, Brian K Kobilka, and Jan Steyaert. A general protocol for the generation of nanobodies for structural biology. Nat. Protoc., 9(3):674–693, March 2014.

12. Iwan Zimmermann, Pascal Egloff, Cedric Aj Hutter, Fabian M Arnold, Peter Stohler, Nicolas Bocquet, Melanie N Hug, Sylwia Huber, Martin Siegrist, Lisa Hetemann, Jennifer Gera, Samira Gmür, Peter Spies, Daniel Gygax, Eric R Geertsma, Roger Jp Dawson, and Markus A Seeger. Synthetic single domain antibodies for the conformational trapping of membrane proteins. Elife, 7, May 2018.

13. Sandrine Moutel, Nicolas Bery, Virginie Bernard, Laura Keller, Emilie Lemesre, Ario de Marco, Laetitia Ligat, Jean-Christophe Rain, Gilles Favre, Aurélien Olichon, and Franck Perez. NaLi-H1: A universal synthetic library of humanized nanobodies providing highly functional antibodies and intrabodies. Elife, 5, July 2016.

14. Florian I Schmidt, Leo Hanke, Benjamin Morin, Rebeccah Brewer, Vesna Brusic, Sean P J Whelan, and Hidde L Ploegh. Phenotypic lentivirus screens to identify functional single domain antibodies. Nat Microbiol, 1(8):16080, June 2016.

15. Enkelejda Miho, Rok Roškar, Victor Greiff, and Sai T Reddy. Large-scale network analysis reveals the sequence space architecture of antibody repertoires. Nat. Commun., 10(1): 1321, March 2019.

16. Hinrich Schütze, Christopher D Manning, and Prabhakar Raghavan. Introduction to information retrieval, volume 39. Cambridge University Press Cambridge, 2008.

17. Etienne Becht, Leland McInnes, John Healy, Charles-Antoine Dutertre, Immanuel W H Kwok, Lai Guan Ng, Florent Ginhoux, and Evan W Newell. Dimensionality reduction for visualizing single-cell data using UMAP. Nat. Biotechnol., December 2018.

18. Venkatesh Kumar, Thomas Vollbrecht, Mark Chernyshev, Sanjay Mohan, Brian Hanst, Nicholas Bavafa, Antonia Lorenzo, Nikesh Kumar, Robert Ketteringham, Kemal Eren, Michael Golden, Michelli F Oliveira, and Ben Murrell. Long-read amplicon denoising. Nucleic Acids Res., 47(18):e104, October 2019.

19. Leo Hanke, Hrishikesh Das, Daniel J Sheward, Laura Perez Vidakovics, Egon Urgard, Ainhoa Moliner-Morro, Changil Kim, Vivien Karl, Alec Pankow, Natalie L Smith, Bartlomiej Porebski, Oscar Fernandez-Capetillo, Erdinc Sezgin, Gabriel K Pedersen, Jonathan M Coquet, B Martin Hällberg, Ben Murrell, and Gerald M McInerney. A bispecific monomeric nanobody induces spike trimer dimers and neutralizes sars-cov-2 in vivo. bioRxiv, 2021. doi: 10.1101/2021.03.20.436243.

20. Erik Volz, Swapnil Mishra, Meera Chand, Jeffrey C Barrett, Robert Johnson, Lily Geidelberg, Wes R Hinsley, Daniel J Laydon, Gavin Dabrera, Áine O’Toole, Roberto Amato, Manon Ragonnet-Cronin, Ian Harrison, Ben Jackson, Cristina V Ariani, Olivia Boyd, Nicholas J Loman, John T McCrone, Sónia Gonçalves, David Jorgensen, Richard Myers, Verity Hill, David K Jackson, Katy Gaythorpe, Natalie Groves, John Sillitoe, Dominic P Kwiatkowski, COVID-19 Genomics UK (COG-UK) consortium, Seth Flaxman, Oliver Ratmann, Samir Bhatt, Susan Hopkins, Axel Gandy, Andrew Rambaut, and Neil M Ferguson. Assessing transmissibility of SARS-CoV-2 lineage b.1.1.7 in england. Nature, pages 1–17, March 2021.

21. Houriiyah Tegally, Eduan Wilkinson, Marta Giovanetti, Arash Iranzadeh, Vagner Fonseca, Jennifer Giandhari, Deelan Doolabh, Sureshnee Pillay, Emmanuel James San, Nokukhanya Msomi, Koleka Mlisana, Anne von Gottberg, Sibongile Walaza, Mushal Allam, Arshad Ismail, Thabo Mohale, Allison J Glass, Susan Engelbrecht, Gert Van Zyl, Wolfgang Preiser, Francesco Petruccione, Alex Sigal, Diana Hardie, Gert Marais, Nei-Yuan Hsiao, Stephen Korsman, Mary-Ann Davies, Lynn Tyers, Innocent Mudau, Denis York, Caroline Maslo, Dominique Goedhals, Shareef Abrahams, Oluwakemi Laguda-Akingba, Arghavan Alisoltani-Dehkordi, Adam Godzik, Constantinos Kurt Wibmer, Bryan Trevor Sewell, José Lourenço, Luiz Carlos Junior Alcantara, Sergei L Kosakovsky Pond, Steven Weaver, Darren Martin, Richard J Lessells, Jinal N Bhiman, Carolyn Williamson, and Tulio de Oliveira. Detection of a SARS-CoV-2 variant of concern in south africa. Nature, 592(7854):438–443, April 2021.

22. Nuno R Faria, Thomas A Mellan, Charles Whittaker, Ingra M Claro, Darlan da S Candido, Swapnil Mishra, Myuki A E Crispim, Flavia C Sales, Iwona Hawryluk, John T McCrone, Ruben J G Hulswit, Lucas A M Franco, Mariana S Ramundo, Jaqueline G de Jesus, Pamela S Andrade, Thais M Coletti, Giulia M Ferreira, Camila A M Silva, Erika R Manuli, Rafael H M Pereira, Pedro S Peixoto, Moritz U Kraemer, Nelson Gaburo, Cecilia da C Camilo, Henrique Hoeltgebaum, William M Souza, Esmenia C Rocha, Leandro M de Souza, Mariana C de Pinho, Leonardo J T Araujo, Frederico S V Malta, Aline B de Lima, Joice do P Silva, Danielle A G Zauli, Alessandro C de S Ferreira, Ricardo P Schnekenberg, Daniel J Laydon, Patrick G T Walker, Hannah M Schlüter, Ana L P Dos Santos, Maria S Vidal, Valentina S Del Caro, Rosinaldo M F Filho, Helem M Dos Santos, Renato S Aguiar, José L P Modena, Bruce Nelson, James A Hay, Melodie Monod, Xenia Miscouridou, Helen Coupland, Raphael Sonabend, Michaela Vollmer, Axel Gandy, Marc A Suchard, Thomas A Bowden, Sergei L K Pond, Chieh-Hsi Wu, Oliver Ratmann, Neil M Ferguson, Christopher Dye, Nick J Loman, Philippe Lemey, Andrew Rambaut, Nelson A Fraiji, Maria do P S S Carvalho, Oliver G Pybus, Seth Flaxman, Samir Bhatt, and Ester C Sabino. Genomics and epidemiology of a novel SARS-CoV-2 lineage in manaus, brazil. medRxiv, March 2021.

23. Constantinos Kurt Wibmer, Frances Ayres, Tandile Hermanus, Mashudu Madzivhandila, Prudence Kgagudi, Brent Oosthuysen, Bronwen E Lambson, Tulio de Oliveira, Marion Vermeulen, Karin van der Berg, Theresa Rossouw, Michael Boswell, Veronica Ueckermann, Susan Meiring, Anne von Gottberg, Cheryl Cohen, Lynn Morris, Jinal N Bhiman, and Penny L Moore. SARS-CoV-2 501Y.V2 escapes neutralization by south african COVID-19 donor plasma. Nat. Med., 27(4):622–625, April 2021.

24. Michael Schoof, Bryan Faust, Reuben A Saunders, Smriti Sangwan, Veronica Rezelj, Nick Hoppe, Morgane Boone, Christian B Billesbølle, Cristina Puchades, Caleigh M Azumaya, Huong T Kratochvil, Marcell Zimanyi, Ishan Deshpande, Jiahao Liang, Sasha Dickinson, Henry C Nguyen, Cynthia M Chio, Gregory E Merz, Michael C Thompson, Devan Diwanji, Kaitlin Schaefer, Aditya A Anand, Niv Dobzinski, Beth Shoshana Zha, Camille R Simoneau, Kristoffer Leon, Kris M White, Un Seng Chio, Meghna Gupta, Mingliang Jin, Fei Li, Yanxin Liu, Kaihua Zhang, David Bulkley, Ming Sun, Amber M Smith, Alexandrea N Rizo, Frank Moss, Axel F Brilot, Sergei Pourmal, Raphael Trenker, Thomas Pospiech, Sayan Gupta, Benjamin Barsi-Rhyne, Vladislav Belyy, Andrew W Barile-Hill, Silke Nock, Yuwei Liu, Nevan J Krogan, Corie Y Ralston, Danielle L Swaney, Adolfo García-Sastre, Melanie Ott, Marco Vignuzzi, QCRG Structural Biology Consortium, Peter Walter, and Aashish Manglik. An ultrapotent synthetic nanobody neutralizes SARS-CoV-2 by stabilizing inactive spike. Science, 370(6523):1473–1479, December 2020.

25. John Jumper, Richard Evans, Alexander Pritzel, Tim Green, Michael Figurnov, Olaf Ronneberger, Kathryn Tunyasuvunakool, Russ Bates, Augustin Žídek, Anna Potapenko, Alex Bridgland, Clemens Meyer, Simon A A Kohl, Andrew J Ballard, Andrew Cowie, Bernardino Romera-Paredes, Stanislav Nikolov, Rishub Jain, Jonas Adler, Trevor Back, Stig Petersen, David Reiman, Ellen Clancy, Michal Zielinski, Martin Steinegger, Michalina Pacholska, Tamas Berghammer, Sebastian Bodenstein, David Silver, Oriol Vinyals, Andrew W Senior, Koray Kavukcuoglu, Pushmeet Kohli, and Demis Hassabis. Highly accurate protein structure prediction with AlphaFold. Nature, July 2021.

26. Sergey Ovchinnikov, Milot Mirdita, and Martin Steinegger. ColabFold - making protein folding accessible to all via google colab, July 2021.

27. Thomas E Wales and John R Engen. Hydrogen exchange mass spectrometry for the analysis of protein dynamics. Mass Spectrom. Rev., 25(1):158–170, January 2006.

28. Z Zhang and D L Smith. Determination of amide hydrogen exchange by mass spectrometry: a new tool for protein structure elucidation. Protein Sci., 2(4):522–531, April 1993.

29. Christopher O Barnes, Claudia A Jette, Morgan E Abernathy, Kim-Marie A Dam, Shannon R Esswein, Harry B Gristick, Andrey G Malyutin, Naima G Sharaf, Kathryn E Huey-Tubman, Yu E Lee, Davide F Robbiani, Michel C Nussenzweig, Anthony P West, Jr, and Pamela J Bjorkman. SARS-CoV-2 neutralizing antibody structures inform therapeutic strategies. Nature, 588(7839):682–687, December 2020.

30. Seth J Zost, Pavlo Gilchuk, James Brett Case, Elad Binshtein, Rita E Chen, Joseph P Nkolola, Alexandra Schäfer, Joseph X Reidy, Andrew Trivette, Rachel S Nargi, Rachel E Sutton, Naveenchandra Suryadevara, David R Martinez, Lauren E Williamson, Elaine C Chen, Taylor Jones, Samuel Day, Luke Myers, Ahmed O Hassan, Natasha M Kafai, Emma S Winkler, Julie M Fox, Swathi Shrihari, Benjamin K Mueller, Jens Meiler, Abishek Chandrashekar, Noe B Mercado, James J Steinhardt, Kuishu Ren, Yueh-Ming Loo, Nicole L Kallewaard, Broc T McCune, Shamus P Keeler, Michael J Holtzman, Dan H Barouch, Lisa E Gralinski, Ralph S Baric, Larissa B Thackray, Michael S Diamond, Robert H Carnahan, and James E Crowe, Jr. Potently neutralizing and protective human antibodies against SARS-CoV-2. Nature, 584(7821):443–449, August 2020.

31. Allison J Greaney, Tyler N Starr, Pavlo Gilchuk, Seth J Zost, Elad Binshtein, Andrea N Loes, Sarah K Hilton, John Huddleston, Rachel Eguia, Katharine H D Crawford, Adam S Dingens, Rachel S Nargi, Rachel E Sutton, Naveenchandra Suryadevara, Paul W Rothlauf, Zhuoming Liu, Sean P J Whelan, Robert H Carnahan, James E Crowe, Jr, and Jesse D Bloom. Complete mapping of mutations to the SARS-CoV-2 spike Receptor-Binding domain that escape antibody recognition. Cell Host Microbe, 29(1):44–57.e9, January 2021.

32. Meng Yuan, Nicholas C Wu, Xueyong Zhu, Chang-Chun D Lee, Ray T Y So, Huibin Lv, Chris K P Mok, and Ian A Wilson. A highly conserved cryptic epitope in the receptor binding domains of SARS-CoV-2 and SARS-CoV. Science, 368(6491):630–633, May 2020.

33. Paul B McCray, Jr, Lecia Pewe, Christine Wohlford-Lenane, Melissa Hickey, Lori Manzel, Lei Shi, Jason Netland, Hong Peng Jia, Carmen Halabi, Curt D Sigmund, David K Meyerholz, Patricia Kirby, Dwight C Look, and Stanley Perlman. Lethal infection of K18-hACE2 mice infected with severe acute respiratory syndrome coronavirus. J. Virol., 81(2):813–821, January 2007.

34. Emma S Winkler, Adam L Bailey, Natasha M Kafai, Sharmila Nair, Broc T McCune, Jinsheng Yu, Julie M Fox, Rita E Chen, James T Earnest, Shamus P Keeler, Jon H Ritter, Liang-I Kang, Sarah Dort, Annette Robichaud, Richard Head, Michael J Holtzman, and Michael S Diamond. SARS-CoV-2 infection of human ACE2-transgenic mice causes severe lung inflammation and impaired function. Nat. Immunol., 21(11):1327–1335, November 2020.

35. Rob C Roovers, Maria J W D Vosjan, Toon Laeremans, Rachid el Khoulati, Renée C G de Bruin, Kathryn M Ferguson, Arie J Verkleij, Guus A M S van Dongen, and Paul M P van Bergen en Henegouwen. A biparatopic anti-EGFR nanobody efficiently inhibits solid tumour growth. Int. J. Cancer, 129(8):2013–2024, October 2011.

36. Ainhoa Moliner-Morro, Daniel J Sheward, Vivien Karl, Laura Perez Vidakovics, Ben Murrell, Gerald M McInerney, and Leo Hanke. Picomolar SARS-CoV-2 neutralization using Multi-Arm PEG nanobody constructs. Biomolecules, 10(12), December 2020.

37. Lorena Itatí Ibañez, Marina De Filette, Anna Hultberg, Theo Verrips, Nigel Temperton, Robin A Weiss, Wesley Vandevelde, Bert Schepens, Peter Vanlandschoot, and Xavier Saelens. Nanobodies with in vitro neutralizing activity protect mice against H5N1 influenza virus infection. J. Infect. Dis., 203(8):1063–1072, April 2011.

38. Ganesh E Phad, Pradeepa Pushparaj, Karen Tran, Viktoriya Dubrovskaya, Monika Àdori, Paola Martinez-Murillo, Néstor Vázquez Bernat, Suruchi Singh, Gilman Dionne, Sijy O’Dell, Komal Bhullar, Sanjana Narang, Chiara Sorini, Eduardo J Villablanca, Christopher Sundling, Benjamin Murrell, John R Mascola, Lawrence Shapiro, Marie Pancera, Marcel Martin, Martin Corcoran, Richard T Wyatt, and Gunilla B Karlsson Hedestam. Extensive dissemination and intraclonal maturation of HIV env vaccine-induced B cell responses, 2020.

39. Kimberly M Cirelli, Diane G Carnathan, Bartek Nogal, Jacob T Martin, Oscar L Rodriguez, Amit A Upadhyay, Chiamaka A Enemuo, Etse H Gebru, Yury Choe, Federico Viviano, Catherine Nakao, Matthias G Pauthner, Samantha Reiss, Christopher A Cottrell, Melissa L Smith, Raiza Bastidas, William Gibson, Amber N Wolabaugh, Mariane B Melo, Benjamin Cossette, Venkatesh Kumar, Nirav B Patel, Talar Tokatlian, Sergey Menis, Daniel W Kulp, Dennis R Burton, Ben Murrell, William R Schief, Steven E Bosinger, Andrew B Ward, Corey T Watson, Guido Silvestri, Darrell J Irvine, and Shane Crotty. Slow delivery immunization enhances HIV neutralizing antibody and germinal center responses via modulation of immunodominance. Cell, 180(1):206, January 2020.

40. Colin Havenar-Daughton, Diane G Carnathan, Archana V Boopathy, Amit A Upadhyay, Ben Murrell, Samantha M Reiss, Chiamaka A Enemuo, Etse H Gebru, Yury Choe, Pallavi Dhadvai, Federico Viviano, Kirti Kaushik, Jinal N Bhiman, Bryan Briney, Dennis R Burton, Steven E Bosinger, William R Schief, Darrell J Irvine, Guido Silvestri, and Shane Crotty. Rapid germinal center and antibody responses in non-human primates after a single nanoparticle vaccine immunization. Cell Rep., 29(7):1756–1766.e8, November 2019.

41. Elise Landais, Ben Murrell, Bryan Briney, Sasha Murrell, Kimmo Rantalainen, Zachary T Berndsen, Alejandra Ramos, Lalinda Wickramasinghe, Melissa Laird Smith, Kemal Eren, Natalia de Val, Mengyu Wu, Audrey Cappelletti, Jeffrey Umotoy, Yolanda Lie, Terri Wrin, Paul Algate, Po-Ying Chan-Hui, Etienne Karita, IAVI Protocol C Investigators, IAVI African HIV Research Network, Andrew B Ward, Ian A Wilson, Dennis R Burton, Davey Smith, Sergei L Kosakovsky Pond, and Pascal Poignard. HIV envelope glycoform heterogeneity and localized diversity govern the initiation and maturation of a V2 apex broadly neutralizing antibody lineage. Immunity, 47(5):990–1003.e9, November 2017.

42. Jeffrey Umotoy, Bernard S Bagaya, Collin Joyce, Torben Schiffner, Sergey Menis, Karen L Saye-Francisco, Trevor Biddle, Sanjay Mohan, Thomas Vollbrecht, Oleksander Kalyuzhniy, Sharon Madzorera, Dale Kitchin, Bronwen Lambson, Molati Nonyane, William Kilembe, IAVI Protocol C Investigators, IAVI African HIV Research Network, Pascal Poignard, William R Schief, Dennis R Burton, Ben Murrell, Penny L Moore, Bryan Briney, Devin Sok, and Elise Landais. Rapid and focused maturation of a VRC01-Class HIV broadly neutralizing antibody lineage involves both binding and accommodation of the N276-Glycan. Immunity, 51(1): 141–154.e6, July 2019.

43. Jessica R Ingram, Kevin E Knockenhauer, Benedikt M Markus, Joseph Mandelbaum, Alexander Ramek, Yibing Shan, David E Shaw, Thomas U Schwartz, Hidde L Ploegh, and Sebastian Lourido. Allosteric activation of apicomplexan calcium-dependent protein kinases. Proc. Natl. Acad. Sci. U. S. A., 112(36):E4975–84, September 2015.

44. Hundeep Kaur, Jean-Baptiste Hartmann, Roman P Jakob, Michael Zahn, Iwan Zimmermann, Timm Maier, Markus A Seeger, and Sebastian Hiller. Identification of conformation-selective nanobodies against the membrane protein insertase BamA by an integrated structural biology approach. J. Biomol. NMR, 73(6-7):375–384, July 2019.

45. Meng Yuan, Hejun Liu, Nicholas C Wu, Chang-Chun D Lee, Xueyong Zhu, Fangzhu Zhao, Deli Huang, Wenli Yu, Yuanzi Hua, Henry Tien, Thomas F Rogers, Elise Landais, Devin Sok, Joseph G Jardine, Dennis R Burton, and Ian A Wilson. Structural basis of a shared antibody response to SARS-CoV-2. Science, 369(6507):1119–1123, August 2020.

46. Christopher O Barnes, Anthony P West, Jr, Kathryn E Huey-Tubman, Magnus A G Hoffmann, Naima G Sharaf, Pauline R Hoffman, Nicholas Koranda, Harry B Gristick, Christian Gaebler, Frauke Muecksch, Julio C Cetrulo Lorenzi, Shlomo Finkin, Thomas Hägglöf, Arlene Hurley, Katrina G Millard, Yiska Weisblum, Fabian Schmidt, Theodora Hatziioannou, Paul D Bieniasz, Marina Caskey, Davide F Robbiani, Michel C Nussenzweig, and Pamela J Bjorkman. Structures of human antibodies bound to SARS-CoV-2 spike reveal common epitopes and recurrent features of antibodies. Cell, 182(4):828–842.e16, August 2020.

47. Agnieszka M Sziemel, Shi-Hsia Hwa, Alex Sigal, Grace Tyson, Nicola Logan, Brian J Willett, and Peter J Durcan. Development of highly potent neutralising nanobodies against multiple SARS-CoV-2 variants including the variant of concern b.1.351.

48. Dapeng Sun, Zhe Sang, Yong Joon Kim, Yufei Xiang, Tomer Cohen, Anna K Belford, Alexis Huet, James F Conway, Ji Sun, Derek J Taylor, Dina Schneidman-Duhovny, Cheng Zhang, Wei Huang, and Yi Shi. Potent neutralizing nanobodies resist convergent circulating variants of SARS-CoV-2 by targeting novel and conserved epitopes. bioRxiv, March 2021.

49. H L Wells, M Letko, G Lasso, B Ssebide, J Nziza, D K Byarugaba, I Navarrete-Macias, E Liang, M Cranfield, B A Han, M W Tingley, M Diuk-Wasser, T Goldstein, C K Johnson, J A K Mazet, K Chandran, V J Munster, K Gilardi, and S J Anthony. The evolutionary history of ACE2 usage within the coronavirus subgenus sarbecovirus. Virus Evol, 7(1):veab007, January 2021.

50. Cécile Vincke, Remy Loris, Dirk Saerens, Sergio Martinez-Rodriguez, Serge Muyldermans, and Katja Conrath. General strategy to humanize a camelid single-domain antibody and identification of a universal humanized nanobody scaffold. J. Biol. Chem., 284(5):3273–3284, January 2009.

51. Daniel Wrapp, Nianshuang Wang, Kizzmekia S Corbett, Jory A Goldsmith, Ching-Lin Hsieh, Olubukola Abiona, Barney S Graham, and Jason S McLellan. Cryo-EM structure of the 2019-nCoV spike in the prefusion conformation. Science, 367(6483):1260–1263, March 2020.

52. Jon Louis Bentley. Multidimensional binary search trees used for associative searching. Commun. ACM, 18(9):509–517, September 1975.

53. Thomas F Rogers, Fangzhu Zhao, Deli Huang, Nathan Beutler, Alison Burns, Wan-Ting He, Oliver Limbo, Chloe Smith, Ge Song, Jordan Woehl, Linlin Yang, Robert K Abbott, Sean Callaghan, Elijah Garcia, Jonathan Hurtado, Mara Parren, Linghang Peng, Sydney Ramirez, James Ricketts, Michael J Ricciardi, Stephen A Rawlings, Nicholas C Wu, Meng Yuan, Davey M Smith, David Nemazee, John R Teijaro, James E Voss, Ian A Wilson, Raiees Andrabi, Bryan Briney, Elise Landais, Devin Sok, Joseph G Jardine, and Dennis R Burton. Isolation of potent SARS-CoV-2 neutralizing antibodies and protection from disease in a small animal model. Science, June 2020.

54. John R Engen and Thomas E Wales. Analytical aspects of hydrogen exchange mass spectrometry. Annu. Rev. Anal. Chem., 8:127–148, May 2015.

55. Victor M Corman, Olfert Landt, Marco Kaiser, Richard Molenkamp, Adam Meijer, Daniel Kw Chu, Tobias Bleicker, Sebastian Brünink, Julia Schneider, Marie Luisa Schmidt, Daphne Gjc Mulders, Bart L Haagmans, Bas van der Veer, Sharon van den Brink, Lisa Wijsman, Gabriel Goderski, Jean-Louis Romette, Joanna Ellis, Maria Zambon, Malik Peiris, Herman Goossens, Chantal Reusken, Marion Pg Koopmans, and Christian Drosten. Detection of 2019 novel coronavirus (2019-nCoV) by real-time RT-PCR. Euro Surveill., 25(3), January 2020.

56. Takayuki Ishige, Shota Murata, Toshibumi Taniguchi, Akiko Miyabe, Kouichi Kitamura, Kenji Kawasaki, Motoi Nishimura, Hidetoshi Igari, and Kazuyuki Matsushita. Highly sensitive detection of SARS-CoV-2 RNA by multiplex rRT-PCR for molecular diagnosis of COVID-19 by clinical laboratories. Clin. Chim. Acta, 507:139–142, August 2020.

57. Roman Wölfel, Victor M Corman, Wolfgang Guggemos, Michael Seilmaier, Sabine Zange, Marcel A Müller, Daniela Niemeyer, Terry C Jones, Patrick Vollmar, Camilla Rothe, Michael Hoelscher, Tobias Bleicker, Sebastian Brünink, Julia Schneider, Rosina Ehmann, Katrin Zwirglmaier, Christian Drosten, and Clemens Wendtner. Virological assessment of hospitalized patients with COVID-2019. Nature, 581(7809):465–469, May 2020.

